# IGS38, a lncRNA from the human rDNA intergenic spacer, regulates rRNA transcription by altering rDNA chromatin organisation and activating the transcription machinery

**DOI:** 10.64898/2026.05.02.722362

**Authors:** Kanwal Tariq, Mareike Polenkowski, Jaclyn Quin, Anaswara Sugathan, Signe Isacson, Stefanie Jakobsson, Elin Enervald, Anne von Euler, Anita Öst, Neus Visa, Ann-Kristin Östlund Farrants

**Author notes:** Corresponding author: Ann-Kristin Östlund Farrants, Department of Molecular Biosciences, The Wenner-Gren Institute, The Arrhenius Lab F4, Stockholm University, SE 106 91 Stockholm, Sweden. Contributed equally to this *work (Kanwal Tariq, Mareike Polenkowski).

## Abstract

The eukaryotic ribosomal genes are multi-copy genes, transcribed from the rDNA, and approximately one third of them is actively transcribed in differentiated cells. A number of lncRNAs have been identified from the intergenic spacer between the rRNA genes, among those the spacer RNA and PAPAS that are involved silencing of rRNA gene copies by altering the chromatin configuration. Here, we have identified lncRNAs that are transcribed from the human rDNA loci and modulate the loci; IGS38 positively regulates rRNA gene transcription by associating to the 47S rRNA gene promoter and modulating the rRNA promoter accessibility while IGS32as associates with heterochromatin. IGS38 binds to the 47S gene promoter through the RNA pol I factors TAF1C and RRN3 as well as the Williams Syndrome Transcription Factor (WSTF), a component of the B-WICH chromatin remodelling complex. The increased accessibility of the promoter stabilises the architectural protein Upstream Binding Factor (UBF) at the rRNA promoter, thereby facilitating RNA pol I promoter escape. Furthermore, IGS38 knock down displays and increased dsRNA abundance in the cytoplasm with a weak induction of the dsRNA sensor OAS2, typically induced by interferon and viral dsRNA. Overall, the both IGS38 and IGS32as are chromatin associated lncRNAs involved in rDNA chromatin changes, and IGS38 is stimulating, together with WSTF, rRNA gene transcription in human cells.

**Graphical abstract:** 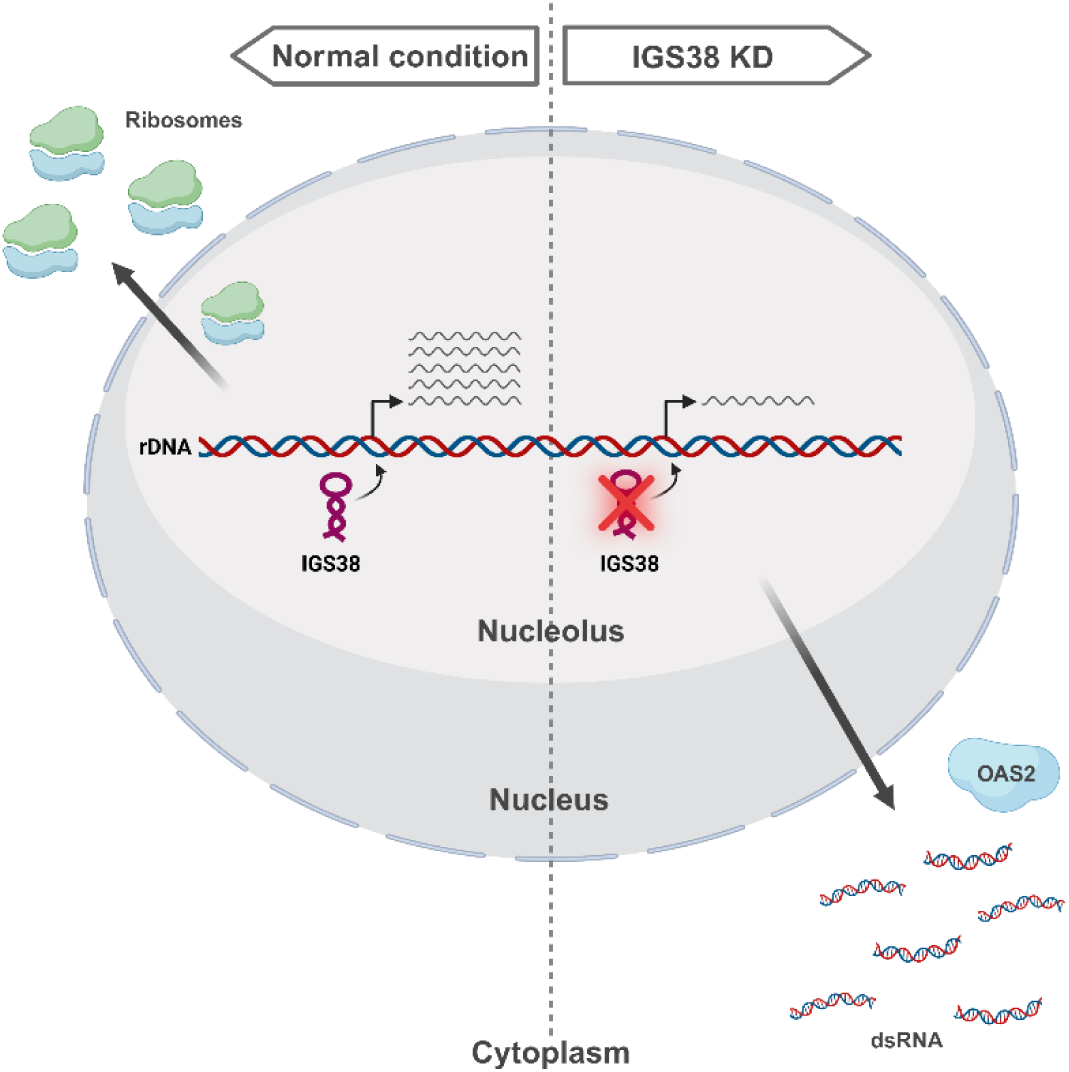

**IGS stabilises 47S rRNA transcription, disruption of IGS38 expression leads to the release of dsRNA in the cytoplasm and a weak immune activation of OAS2.**

Created by biorender (https://biorender.com/shortURL)

## Introduction

Ribosomal RNA synthesis in the nucleolus is the first step in ribosomal biogenesis, in which ribosomes are assembled for protein synthesis. The nucleolar 47S rRNA gene transcription is highly regulated, with modifications of the specific RNA polymerase I (RNA pol I) machinery being essential, but also factors such as chromatin remodelling complexes, histone and DNA modifying enzymes, and noncoding RNAs play an important role. In particular, maintaining an accessible chromatin state at the promoter as well as in the transcribed region is necessary for efficient transcription (reviewed in 1-3). The 47S rRNA transcript is transcribed from the ribosomal rRNA gene repeats arranged as tandem repeats in the rDNA loci at acrocentric chromosomes (1–3). The diploid human genome has hundreds of these gene repeats but only half to a third of these repeats are actively transcribed and the others are maintained in inactive or silent states in differentiated cells (2–6).

In addition to the 47S rRNA genes, several noncoding RNAs originate from the rDNA intergenic spacer (IGS) region and these are associated with different functions (reviewed in 6-10). Two transcripts close to the 47S rRNA promoter affect the 47S rRNA gene transcription; the spacer-RNA harbouring the pRNA (promoter RNA) and PAPAS (promoter and pre-rRNA antisense) transcripts was first identified in mice, are involved in transcriptional silencing by recruiting chromatin remodelling complexes. The spacer-RNA is transcribed from the upstream spacer promoter (5, 11–13) and generates the 250bp pRNA fragment which forms RNA:DNA triplex hybrids at the rRNA promoter (14–16). This structure, together with the TTF1 (Terminator Transcription terminator factor 1) bound at the promoter, recruits TIP5 (TTF1 interacting protein 5) in NoRC (nucleolar remodelling complex) to establish a silent chromatin state with DNA methylation and inactivating histone modifications, including H3K9me3 and HP1 (heterochromatin protein 1) (16–20). The antisense PAPAS transcripts, encompassing the rRNA gene promoter, also form RNA:DNA hybrids with the promoter (21–24). PAPAS recruit the histone methyltransferase Suv4-20h2 when cells become quiescent (25) and the NuRD (Nucleosome Remodelling Deacetylase) complex to the promoter in response to heat shock and hypotonic stress (22, 24, 26). The chromatin silencing includes moving a nucleosome over TSS at the promoter and, establishing a so-called poised state, to prevent factor recruitment and thus reduces transcription (21, 22). The poised chromatin state is activated by chromatin remodelling complexes CSB (Cockayne syndrome group B), which move the nucleosome from the TSS and establishes an active acetylated chromatin state (21, 27, 28). Activation of rRNA 47S gene transcription is mainly regulated by modifications of the RNA pol I machinery, such as phosphorylation of RRN3/TIF-1A (RNA polymerase-I transcription initiation factor 1A) and UBF (1, 2), but it also involves alterations of the chromatin at the promotor by chromatin remodelling complexes, such as CSB and B-WICH, which alters the accessibility at the 47S rRNA promoter (29–32).

Further ncRNAs have been identified from the human IGS, which in contrast to the rRNA gene body is not evolutionary conserved, even between mammals. Counterparts to the mouse pRNA and PAPAS have been identified in human cell, both associated with R-loops, which increase when not resolved by RPA (replication protein A), Senataxin, RNAse H and Topoisomerase (33–35). The nucleolus is a stress hub, sensing different stressors, such as viral infections, nutrient starvation, heat shock and hypotonic stress, often resulting in p53 induced nucleolar stress leading to apoptosis (36, 37). Nucleolar stress also induces p53 independent responses through ribosomal proteins, HIF1a and the immune response signalling molecule NFκB (37, 38). pRNA and PAPAS are activated upon stress and dysregulated in cancer cells, and expression of PAPAS is associated with an induced immune response, with a number of interferon stimulated genes, such as Irf7, being expressed (35). The human PAPAS is initiated by RNA pol II from the 3’ end of the gene repeats and the read through over the promoter is translated into a protein, RIEP (34). This protein is found in the nucleolus and mitochondria upon heat shock (33). Furthermore, a number of short stress induced ncRNA originating from the human IGS have been identified (39, 40). These ncRNA are produced both in an anti-sense direction by RNA pol II, asincRNA (antisense intergenic ncRNA), and in the sense direction, sincRNA (sense intergenic ncRNA), by RNA pol I, with asincRNA maintaining nucleolar structure by forming an R-loop shield (40). This region, from 18 kb to 28 kb from the transcription start site also sequester proteins important in different stress responses, such as VHL and HSP70 (39). This shows that in human cells several ncRNAs contribute to the stress response in the nucleolus.

Stress responses regulating the 47S rRNA gene transcription in human cells also involve lncRNA stemming from outside of the rDNA. RNA pol II transcribed Alu RNAs promote nucleolar assembly and overexpression leads to an increase in nucleolar size and in enhanced 47S rRNA transcription (41). The nucleolar distal regions also produce nucleolar ncRNAs, Disnor RNAs, important for nucleolar structure and 47S rRNA gene transcription by anchoring the rRNA genes to the nucleolar periphery (42–44). Only a few cancer-related lncRNAs, which translocate from their site of origin in the nucleoplasm, actively induce 47S gene transcription by affecting the promoters. LINC01116 RNA associates with rRNA transcriptional upregulation in many cancer cell, such as lung adenocarcinoma, and it interacts with the RNA pol I SL1 complex subunits TAF1A and TAF1D (45). Similarly, SLERT activates rRNA gene transcription by binding to the RNA binding protein DDX21 and releasing its inhibition (46–47).

Here, we have identified ncRNAs originating from the human IGS, one of which, IGS38, had an impact on rRNA transcription. The RNA pol II transcript IGS38 associates with the 47S rRNA gene promoter by binding to the RNA pol I factors TAF1C and RRN3/TIF1A and stabilises the RNA pol I for transcription initiation. The IGS38 also binds to the WSTF (Williams syndrome transcription factor) in the chromatin remodelling complex B-WICH (29, 30), a positive regulator of rRNA gene transcription. This suggests that IGS38 interacts with B-WICH to establish an accessible chromatin configuration at the promoter which then allows factor recruitment and assembly of the RNA pol I machinery. Knock down of IGS38 does not result in an increased transcription of spacer-RNA or PAPAS showing that these RNAs work independently to regulate 47S rRNA transcription. Nevertheless, knock down leads to a weak increase of dsRNA in cytoplasm, mildly inducing the dsRNA sensor OAS2, and Interferon Stimulated Genes (ISG), showing that dysregulation of rRNA leads to an endogenous immune response.

## Methods

### Cell culture and materials

HeLa cells and U2OS cells were maintained in 5% CO2 at 37°C in high glucose DMEM supplemented with 10% FBS (Gibco, Thermo Fisher) and 1% penicillin-streptomycin (Gibco, Thermo Fisher). MCF10 cells (ATCC, US) were maintained in 5% CO2 at 37°C in high glucose DMEM supplemented with 10% FBS (Gibco, Thermo Fisher) with 1 μM insulin and 1% penicillin-streptomycin. Dermal human fibroblasts (ATCC, US) were maintained in 5% CO2 at 37°C in high glucose DMEM supplemented with 10% FBS (Gibco, Thermo Fisher) and 1% penicillin-streptomycin (Gibco, Thermo Fisher).

Peripheral blood cells were retrieved from anonymous, healthy blood donors (Stockholm Blood Centre) who cannot be traced back; thus, the project does not require approval from the Swedish ethical review authority. Primary monocytes and monocyte derived monocytes were isolated from peripheral blood mononuclear cells (PBMC) prepared from buffy coats using Ficoll-Hypaque (Cytiva) gradient centrifugation. Monocytes were enriched from PBMC by negative selection using EasySep^TM^ human monocyte kit (STEMCELL Technologies) according to the manufactureŕs instructions. Monocytes were resuspended at 1×10^6^ cells/ml and were seeded in 96-well plates (with 200 μl/well) or 6-well plates (with 2 ml/well) using RPMI-1640 culture medium which was supplemented with 20 mM HEPES, 2 mM L-glutamine, 100 U/ml penicillin, 100 μg/ml streptomycin (all from Cytiva), 5% human AB serum (Sigma-Aldrich), 50 μM 2-mercaptoethanol, 2% sodium pyruvate (both from Gibco), 35 ng/ml IL-4 and 50 ng/ml GM-CSF (both from PeproTech).

### Transfection of cells

LNA-DNA Gapmer/SiRNA transfections were performed in antibiotic free medium using RNAiMax (Thermo Fisher) transfection reagent. Pre-designed antisense LNA-DNA Gapmers against target and control were from Qiagen, and used at a final concentration of 10nM for all experiments except where noted. Changes in expression were detected 24 hours post transfection. For WSTF knock down, a combination of two target siRNA were used at a concentration of 30 nM for 30 hours. Primer, oligo and probe sequences are listed in Supplementary Table 1. Ribosomal assembly U13369.1 was used as a reference for designing sequences.

Plasmid vector transfections were performed using Lipofectamine 2000 according to the manufacturer’s protocol. Either 2 µg of Empty Vector Control or IGS38 (3 different vectors) cloned with snoRNA Box A and Box B for localization into the nucleolus was used. The sequences for the snoRNA boxes used: box C (5’-TGCAATGATGTCGTAATTTGCGTC-3’) and box D (5’- GACGAAAATTCTTACTGAGCAA -3’) (35). After 48 h of incubation, RNA isolation, cDNA synthesis and qRT-PCR were performed as described below.

### Inhibitor treatments

Cells were treated in antibiotic free medium with 5nM (low dose) or 4μM (high dose) Actinomycin D for 2 hours, or 5 μM α-amanitin or 0.1 μM CX-5461 for 24 hours. RNA was extracted and qPCR was performed as described below.

### Stress treatments

For heat shock, cells were kept at 42°C for 1 hour and subsequently for 1 hour at 37°C before collection. Hypotonic stress was induced by growing cells in media consisting of 30% DMEM (supplemented with FBS as usual but without antibiotics) and 70% H2O. All cells for control treatments were grown in 37°C for the entire treatment time, in 10% FBS supplemented DMEM without antibiotics. Growth factor depletion was performed by growing cells in 1% serum for 48 hours.

### RNA extraction and qPCR

RNA was extracted using Tri-reagent (Thermo Fisher) and converted to cDNA using Superscript III with random primers (Thermo Fisher). RT-qPCRs were analysed using primers against 18S rRNA as internal control with 2^-ΔΔCt^ method. qPCR was performed with KAPA-mix (Merck) or qPCRBIO SYGreen Mix Separate-ROX (PCRBIO).

### Northern blot

Total RNA was extracted by Trizol (Thermo Fisher) and 15μg was loaded on 1% agarose gel. The RNA was probed with specific radiolabelled oligonucleotides after blotting on a Zeta probe membrane (Biorad). Hybridization was performed in ULTRA-hyb oligo buffer at 65°C. Membranes were developed with Fuji Phosphoimager. Probe sequences are listed in the Supplementary Table 1.

### Immunoprecipitation (ChIP and ChRIP)

ChIPs were performed as previously described (31). Briefly, chromatin was cross-linked with 1% formaldehyde for 10 minutes, lysed with buffer (50mM HEPES pH 7.6, 140mM NaCl, 1mM EDTA, 10% glycerol, 0.5% NP-40, and 0.25% Triton X-100). Nuclei were extracted with buffer (200mM NaCl, 1mM EDTA, 0.5mM EGTA, and 10mM Tris pH 8.0) and sonicated in buffer (1mM EDTA, 0.5mM EGTA, and 10mM Tris pH 8.0) in a Bioruptor Plus sonicator 30 seconds high intensity and 30 second paus for 45 minutes. IP was performed with precleared chromatin using target antibodies (Supplementary Table 2). A mixture Protein A/G Sepharose beads were used for capture, followed by reverse crosslinking overnight at 65°C in the presence of proteinase K and RNAse A (Thermo Fisher). DNA was extracted with phenol/chloroform/isoamylalcohol. The samples were analysed qPCR results were calculated as percentage of input after subtracting background using an average of no-antibody and IgG controls. Knock down cells were collected 24 hpt.

ChRIP were performed in a similar manner by modifying the protocol Subhash et al (48), except samples were treated with DNAse after reverse-crosslinking and RNA was extracted with Trizol. After another round of DNAse treatment, results were analysed by qPCR as described above.

### ATAC-qPCR (Assay for Transposase-Accessible Chromatin)

Cells were transfected with control or target LNA-DNA Gapmers for 24 hours and collected following trypsinisation. A total of 25,000 cells were used per treatment, washed with PBS, and resuspended in cold lysis buffer as previously described (49). For tagmentation, we used Illumina Tagment DNA Enzyme and Buffer kit (Illumina; Cat no. 20034197). Tagmentation reactions were incubated at 37°C for 30 minutes and DNA was purified with MinElute (Qiagen) columns. DNA was amplified using a region-specific reverse primer and a tagmentation specific forward primer (49, 50). The 27kb IGS region was used as an internal control.

### CHART-qPCR (Capture hybridization analysis of RNA targets)

Four overlapping target probes were designed in the same region as the probes used for Northern blot and the DNA probes were tested with RNAseH assay as previously described (51), the primer used presented in Supplementary Figure S2B. Briefly, cells were crosslinked with 1% formaldehyde and homogenized. Nuclei were extracted with sucrose and pelleted through a glycerol cushion. Nuclei were sonicated and the extract was cleared with centrifugation. The RNAse H treatment was performed according to the protocol and RNA was extracted with PureLink RNA isolation kit (Thermo Fisher). cDNA was converted using Superscript IV VILO kit (Thermo Fisher). Probes were selected if the target probe reduced RNA abundance but its antisense did not reduce RNA levels according to RNAse H sensitivity analysis.

Target probes with a C18 spacer and 3’biotin tags were from Eurofins Genomics. Hybridization reactions were conducted separately with target probes for IGS38 and IGS32as, and a no probe control for up to 12 hours. Probe bound DNA was captured using Streptavidin labelled MyOne Dynabeads (Thermo Fisher) and treated with RNAseH before eluting. Results were analysed with qPCR using primers against specified regions and presented as fold enrichment compared to input DNA.

### RNA-FISH (Fluorescence in situ hybridization)

Cells were seeded on poly-L-lysine coated slides, fixed with 3.6% formaldehyde and permeabilized with 0.5 % TritonX-100. Cells were washed with 2x saline sodium citrate (SSC) and incubated overnight with target or control probes. Slides were sequentially washed with 50% formamide in 2xSSC, 0.5xSSC, 0.5xSSc in 0.1% Tween 20, and finally with PBS. After blocking with 3% BSA, cells were first incubated in FITC-anti-DIG and anti-Fibrillarin, then incubated with Alexa-anti-FlTC and GAR-Alexa-594. Finally, cells were mounted with Vectashield. Probes were designed with Stellaris Probe Designer (Biosearch Technologies by LGC) and labelled with digoxigenin (DIG) (Eurofins Genomics). The probe used for detection of IGS38 was a probe set including two oligonucleotides corresponding to sequences at position 39.138 kb and 39.152 kb in the rDNA, respectively. Probe sequences are provided in the Supplementary Table 1. Images were acquired with LSM800 laser confocal imager (Zeiss) with ZEN blue edition v3.7.97.07000 software, equipped with a x63 Plan-apochromat 1.4 NA oil-immersion lens (Zeiss). Images were analysed and processed with Fiji/ImageJ2 v14.0/1.53o. The images shown are Z-projections obtained by using maximum intensity projection. Images within one experiment were acquired and processed the same way, unless else is written.

### Dotblot

HeLa cells were transfected with Control, IGS32as or IGS38 antisense LNA-DNA Gapmers. After 30 hours, RNA was harvested as previously described. The RNA was spotted onto a Hybond-N+ membrane (Amersham Biosciences) and left to air dry. The membrane was baked at 80 °C for 2 h and blocked with 5 % milk in PBS-T for 30 min. Primary antibody incubation was performed overnight at 4 °C followed by washes in PBS containing 0.05 % Tween-20 (PBS-T), the secondary antibody (HRP) was added for 1 h at room temperature, washed in PBS-T and developed with Clarity Western ECL Substrate (BioRad). Antibodies used are presented in Supplementary Table 2.

### Immunofluorescence staining

HeLa cells were transfected with LNA-DNA Gapmer-Control or -IGS38 on glass coverslips for 30 hours. Cells were washed with PBS, fixed in 4 % methanol-free Paraformaldehyde for 10 min and permeabilized with PBS containing 0.2 % Triton X-100 for 15 min. Blocking was performed in 5 % BSA containing 0.2 % Triton X-100 for 30 min and the primary antibody for detection of dsRNA (J2, Cell Signaling, 76651) was incubated overnight at 4 °C in 1 % BSA. The cells were washed with 0.1 % Tween-20 in PBS followed by incubation with the secondary antibody (Invitrogen, AlexaFluor 594 (A-11005) diluted in blocking buffer (1:600) for 1 h at room temperature. The cells were washed with PBS before incubation with FITC-phalloidin (Sigma) at a dilution of 1:200 for 30 minutes, before mounted with DAPI containing mounting media (Vectashield, Vector Laboratories) on microscope slides. The slides were viewed with a LSM 800 Airyscan.

### Bioanalyzer analysis

HeLa cells were transfected with 30nM Control or IGS38 antisense LNA-DNA. Total RNA was isolated with the NucleoSpin RNA isolation kit (Macherey-Nagel) and RNA quantification was performed with the Qubit RNA HS Assay Kit (Invitrogen). 5µg/µl of the samples were loaded onto an Agilent high-sensitivity RNA 6000 Pico-Chip and analysed in a 2100 Bioanalyzer system.

### RNA sequencing

Small RNA-seq was performed by lysing freshly frozen HeLa cells with Tissue Lyser LT (Qiagen) with 5mm Stainless Steel Beads (Qiagen) in Qiazol (Qiagen). RNA was extracted with miRNeasy Micro kit (Qiagen). Bioanalyzer (Agilent Technologies, Santa Clara, USA) was used to check RNA integrity, obtaining RIN values between 9.3-10. Library was prepared with NEBNext Small RNA Library Prep Set for Illumina (New England Biolabs, Ipswich, USA) using 100 ng of input RNA. Library preparation was performed according to manufacturer’s instruction with the exception of diluting adapters 1:2 and performing amplification for 12 cycles. Library concentration was determined with QuantiFluor ONE ds DNAsystem on a Quantus fluorometer (Promega, Madison, USA) before pooling. Size selection of libraries was done by running on 6% polyacrylamide Novex TBE gel (Invitrogen, Waltham, USA), where bands of 130-190nt were extracted with Gel breaker tubes (IST Engineering, Milpitas, USA) in buffer provided in the NEBNext kit. The size selected pooled library concentration was determined again before sequencing on NextSeq 500 with NextSeq 500/550 High Output Kit version 2.5, 75 cycles (Illumina, San Diego, USA). Adaptor trimming and sequence count was performed with Seqpac ver. 1.2.0 in R ver. 4.3.2. Mean sequencing depths of all samples were 5,215,570 with a standard deviation of 667,381. Baseline filtering was performed for sequences of size 14 to 75 with a threshold of 5 sequences in 35% of samples prior to counts per million (cpm) normalization. Sequences were mapped to human ribosomal DNA complete repeating unit, GenBank ID:U13369.1. Raw and processed data files are available from GEO with accession number GSE279932.

Transcriptomic sequencing was performed on cell transfected with control scramble LNA-DNA Gapmers or cells transfected with IGS38 target LNA-DNA Gapmers after 24 hpt at a concentration of 30nM, using Eurofins Genomics service. Briefly, reads generated from rRNA sequences were detected using RiboDetector and removed. Poor quality peaks were removed using fastp, also reads shorter than 30bp were removed. The data was then processed in house using Galaxy workflow (www.usegalaxy.org) for subsequent analysis. Bam-files were uploaded to Galaxy and the function FeatureCount (Galaxy Version 2.0.3+galaxy2) was applied with default parameters (paired-end, count as 1 single fragment; gene annotation file: T2T-CHM13v2.0) for read counting. Afterwards, DESeq2 (Galaxy Version 2.11.40.8+galaxy0) was used to determine differentially expressed genes (default parameters, Factor 1: IGS38 KD; Factor 2: control Gapmer). For visualization of the data, a volcano plot (Galaxy Version 0.0.6) was created with labels of the 10 most significant genes. Gene enrichment analysis was performed using online gene ontology tool by GO Consortium (52, 53). Only results with FDR p-value less than 0.05 were considered significant. Raw and processed data files are available from GEO with accession number GSE279931.

### Data Availability

Small RNA sequencing and Secpac analysis: Raw and processed data files from are available from GEO with accession number GSE279932.

RNA seq of ribodepleted total RNA-seq: Raw and processed data files are available from GEO with accession number GSE279931.

**Supplementary Data are available**

## Results

### RNA pol I and RNA pol II both generate novel functional IGS RNAs

Several ncRNAs originating from the IGS region of the human rDNA loci have been identified and two of those, pRNA and PAPAS first identified in mice, down-regulate 47S rRNA gene transcription. To further study ncRNA from the human rDNA, we amplified IGS transcripts using region-specific primers in HeLa cells. Several RNAs were found to originate from a number of positions of the IGS (Supplementary Figure 1A). Three transcripts also appeared as longer transcripts in Northern blots; IGS19as RNA (approximately 500 bp) and IGS32as RNA (approximately 800 bp) in the antisense orientation and IGS38 RNA (approximately 1300 bp) in the sense orientation with respect to 47S rRNA transcription (Figure 1A, depicted on to of Figure 1B). These transcripts aligned to regions where we also found transcription with region specific primers, in particular in the regions 18.5 kb to 23.3 kb and 38.5 kb to 39.5 kb from the TSS (transcription start site). ncRNA have previously been identified from the human rDNA loci and these are found as stable short RNAs, reaching 300 bp, suggested to be fragments or short transcripts (14, 23, 28). To examine whether we could detect short ncRNAs from the human IGS, we performed a small RNA sequencing (sRNA-seq). We found short fragments aligning to the regions around 17 kb and 18 kb from the TSS, around the 37 kb and 39 kb regions (around 37 200 bp and 39 000 bp) and at 42.3 kb close to the TSS (Figure 1B, the transcripts depicted on top). The abundance of fragments around 18 kb and 38 kb from the TSS suggests that the IGS19as and IGS38 were degraded and some of those fragments were stable. In contrast, no small fragments were detected from the 32 kb region. In addition, transcripts aligned to the region -706 to -657 bp upstream of the 47S rRNA TSS (approximately position 42.2 kb) (Figure 1B), covering the previously mapped spacer promotor in human cells (approximately -698 ± 11 bp upstream of the TSS) (13, 54). These spacer transcripts typically ranged from 15-22 nucleotides with the largest fragment of 49 bp, stopping prior to the Tsp site (TTF1 spacer binding site) which binds TTF1 and possibly blocked further transcription elongation.

**Figure 1.**
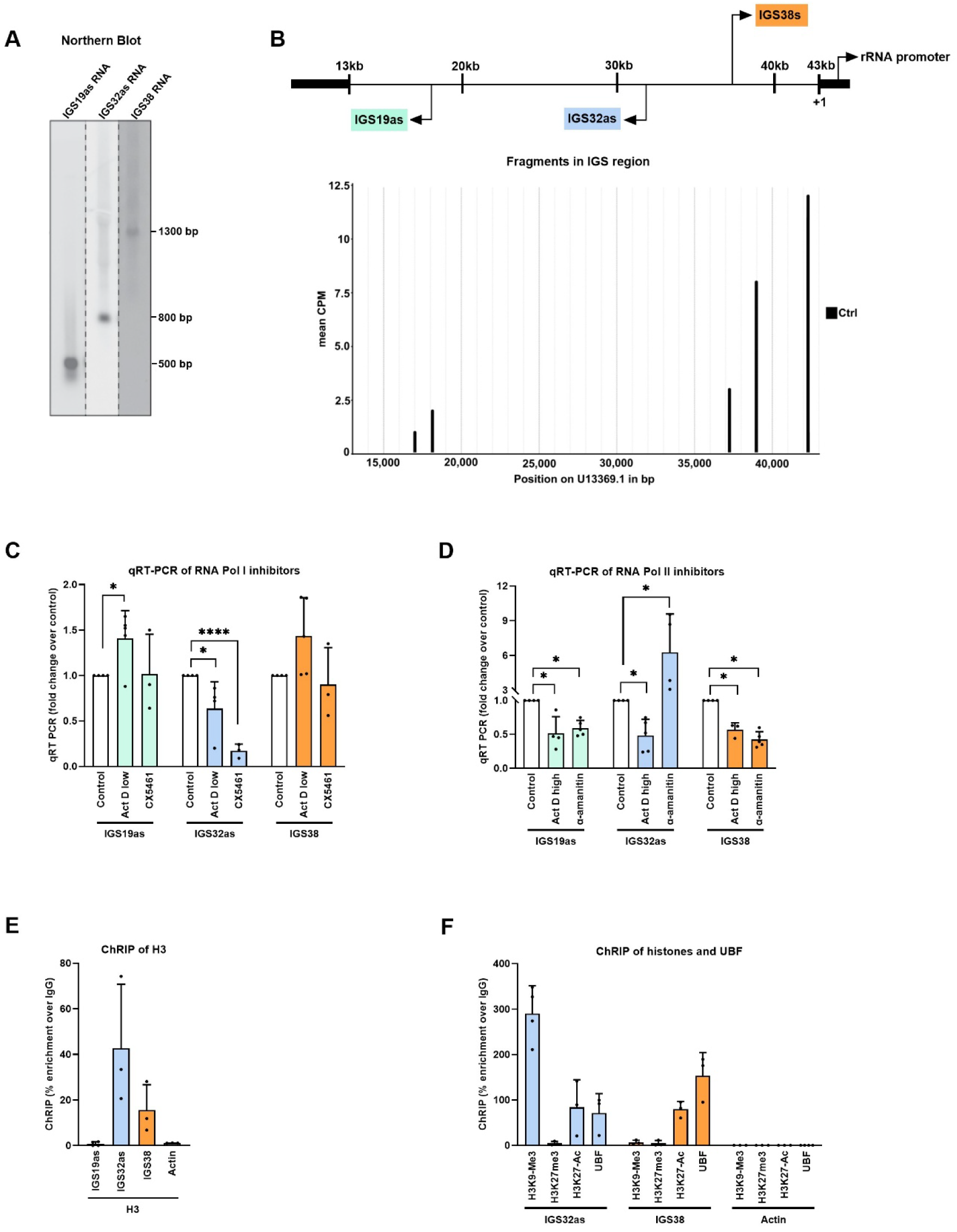
Novel ncRNA from the rDNA are RNA Pol I and RNA Pol II transcripts. A) Identification of stable IGS transcripts by Northern blot. The location of the oligonucleotide probe used for hybridization are indicated at the top of the image. B) Scheme for the positions of the IGS transcripts at the human rDNA gene (top) and the fragments identified by small RNA-seq in control cells (bottom). C) Inhibition of RNA pol I with 5 nM Act D for 2 hours or 0.1 μM CX-5461 for 24 hours, cDNA from total RNA was analysed by qRT-PCR and the signals were normalised to 18S rRNA. The ncRNA detected is marked under the bars. D) Inhibition of RNA pol II with 4 μM Act D for 2 hours or 5 μM α-amanitin for 24 hours, cDNA from total RNA was analysed by qRT-PCR and the signals were normalised to 18S rRNA. The detected ncRNA is marked under the bars. E) ChRIP analysis of the association of IGS19as, IGS32as and IGS38 to chromatin using antibodies against histone H3, the detected RNA is marked under the bars. Actin mRNA is used as a negative control. qRT-PCR data is shown as percentage enrichment over IgG. F) ChRIP analysis of the association of IGS32as and IGS38 to chromatin with histone modifications using antibodies against H3K9me2/3, H3K27me3, H3K27Ac, and UBF, as indicated. Actin mRNA is used as a negative control. qRT-PCR data is shown as percentage enrichment over IgG.= All experiments present means of at least 3 biological replicates. Error bars show the standard deviation. P-values are calculated with unpaired student’s t-test: * p≤ 0.05, **p≤ 0.01,*** p≤ 0.001 and **** p≤ 0.0001.

Since both RNA pol I and RNA pol II have been associated with the human rDNA IGS (25, 33–35, 39, 40) we next examined the occupancy of the polymerases and found both polymerases enriched at the location of the identified lncRNAs (Supplementary Figure S1B). RNA pol II, also in its transcriptionally engaged states with phosphorylation of serin 5 and serin 2 of the carboxyl-terminal domain (CTD), Ser5-P-CTD RNA pol II and Ser2-P-CTD RNA pol II, has been found in a peak around the 38 kb location (39). Using specific antibodies to the Ser5-P-CTD and Ser2-P-CTD RNA pol II, we detected both forms at the 38 kb location (Supplementary Figure S1C), suggesting that this region is actively transcribed by RNA pol II. To elucidate which polymerase that was responsible for the transcription of the lncRNA, we used different RNA polymerase inhibitors; CX-5461 and low concentration of actinomycin D (Act D) were used to inhibit RNA pol I transcription and α-amanitin and Act D in higher concentrations were used for RNA pol II. IGS32as was reduced by RNA pol I inhibitors, in particular CX-5461, while both IGS19as and IGS38 were reduced upon RNA pol II inhibition (Figures 1C and 1D). We conclude that IGS19as and IGS38 are transcribed by RNA pol II and IGS32as is transcribed by RNA pol I. Furthermore, the location and direction of the IGS19as suggests that it is an RNA pol II transcribed asincRNA, preventing RNA pol I aberrant transcription from the region (39, 40).

### The IGS32as associates with constitutive heterochromatin

Many lncRNA are associated with chromatin (55) and the spacer-RNA and PAPAS from the rDNA are associated with promoter chromatin to silence transcription (6, 9, 10). This prompted us to investigate the interactions with chromatin of our identified ncRNAs from the IGS and we performed formaldehyde fixed chromatin-RNA IPs (ChRIP), using antibodies against histone H3. Both IGS32as and IGS38 displayed co-localisation with histone H3-chromatin, whereas IGS19as did not associate to chromatin, similar to the actin mRNA (Figure 1E). The lack of chromatin association of IGS19as further supports that it is an RNA pol II asincRNA (39, 40). We further investigated whether the chromatin-associated RNAs IGS32as and IGS38 associated with different chromatin states by using antibodies against H3K9me3, H3K27me3, H3K27Ac as well as UBF. IGS32as associated with the constitutive heterochromatin mark H3K9me3 but not with the facultative chromatin mark H3K27me3 (Figure 1F). In line with this association, the IGS32as associated with chromatin containing heterochromatin proteins HP1α, but not with EZH2 in PRC2 (Polycomb Repressive Complex 2) (Supplementary Figure S1D), This indicates that IGS332as is associated with constitutive heterochromatin, although associations with active chromatin, H3K27Ac, and UBF were also detected (Figure 1F). IGS38 associated with active chromatin carrying the H3K27Ac mark and with UBF-chromatin, but not with the heterochromatic histone marks (Figure 1F), suggesting that it was associated with active chromatin. IGS38 was not found associated the HP1α and EZH2 at the chromatin (Supplementary Figure S1D). To assess whether these proteins bound directly to the RNAs, we performed RNA immunoprecipitation (RIP) and HP1α bound IGS32as directly, whereas EZH2 did not (Supplementary Figure S1E), further supporting that IGS32as is involved in constitutive heterochromatin. In contrast, IGS38 bound at low levels to the proteins HP1α and EZH2 (Supplementary Figure S1E).

Chromatin-associated RNAs regulate heterochromatin and nuclear architecture, in particular the facultative Polycomb complex binds lncRNAs (55). Since we could not detect interactions between IGS32as and Polycomb we assessed the interaction with constitutive heterochromatin. Heterochromatin associated with H3K9me2/3 and HP1 also depends on lncRNA but different mechanisms may be applied in different species: fission yeast and *Drosophila melanogaster* mainly use RNAi mechanisms (55), but also a mechanism through the RNA exosome is applied by *Drosophila* and mammalian cells (49–57). We knocked down the EXOSC10 component by siRNA and detected an increased level of IGS32as and IGS38, suggesting that they are degraded by the RNA exosome (Supplementary Figure S1F). ChIP performed on EXOSC10 knock down cells showed that an elevated level of HP1α was enriched at the 32kb location compared to control cells (Supplementary Figure S1G), suggesting a similar mechanism to transposon silencing where the presence of nascent RNA is required for HP1α binding (58, 59). To investigate the regulation of IGS32as, we show that it is similarly to the 47S rRNA gene transcription, as glucose stimulation had a tendency to enhance a higher level of IGS32as, but had no effect on the other transcripts (Supplementary Figure S1H). IGS32as transcription was also reduced by reduced activity of c-MYC, a strong regulator of the 47S rRNA transcription (60, 61) and tightly associated with growth, proliferation and metabolic change (62). Knock down of c-MYC by siRNA or by treatment with the c-MYC-MAX inhibitor 10058-F4 reduced the expression of IGS32as (Supplementary Figure S1I), similar to the expression of 47S rRNA (32). We conclude that IGS32as is a heterochromatin RNA, which is regulated in a similar manner as the 47S rRNA transcript.

### IGS38 regulates 47S gene expression and IGS38 knock down changes the chromatin compaction at the 47S rRNA gene promoter

Because of the association of IGS38 RNA with active chromatin, in particular UBF-associated chromatin, we predicted that this ncRNA regulated the 47S rRNA gene transcription. Hence, we knocked down the IGS38 by antisense LNA-DNA Gapmers selected from unique sequences between 38.8 kb and 39.1 kb of this highly repetitive region (sequences in Supplementary Table 1). The transfection of the LNA-DNA Gapmers against IGS38 led to a reduction of the 47S rRNA gene expression in a concentration dependent manner (Figure 2A). In contrast, knock down of IGS32as did not lead to a reduction of the 47S rRNA (Figure 2B). We next expressed fragments from the 38 kb core regions exogenously by transiently transfected RNA pol II expression plasmids. We used three plasmids expressing fragments of 400 bp from 38.759 to 39.156 kb; one fragment covering 200 bp from 38.759 to 38.957 kb, a second one covering 200 bp from 38.958 to 39.156 kb and the third one covering the full 400 bp region, with snoRNAs flanking the fragment (sequences given in Supplementary Table 1). The RNA fragment covering 38.958 to 39.156 kb gave a four-fold increase of the 47S rRNA over controls transfected with empty vector, and the other two RNA fragments gave a slightly lower increase, approximately three-fold (Figure 2C). The expression levels of the IGS38 fragments were approximately similar (Supplementary Figure S2A). As the core fragment of the IGS38 regulated 47S rRNA gene transcription, we next investigated whether IGS38 played a role in changing chromatin states. We designed specific reverse primers and a tagmentation specific forward primer to the promoter and enhancer region (depicted in Figure 2D) to perform ATAC-qPCR. In IGS38 knock down cells, we detected a decrease in chromatin accessibility at the 47S rRNA gene promoter region and an increase at the spacer promoter (Figure 2D). The decrease in accessibility coincided with the TTF1 binding site (T0), UCE and the core promoter (Figure 2E), which is the same region made more accessible by the B-WICH complex (31, 32). IGS32as knock down did not change the accessibility at the 47S rRNA gene promoter, but in contrast to IGS38 knock down led to a less accessible spacer promoter (Figure 2D). Moreover, knock down of IGS38 and IGS32as increased the chromatin compaction at the 32kb region, while neither had an effect on the 38kb region (Supplementary Figure S2D).

**Figure 2.**
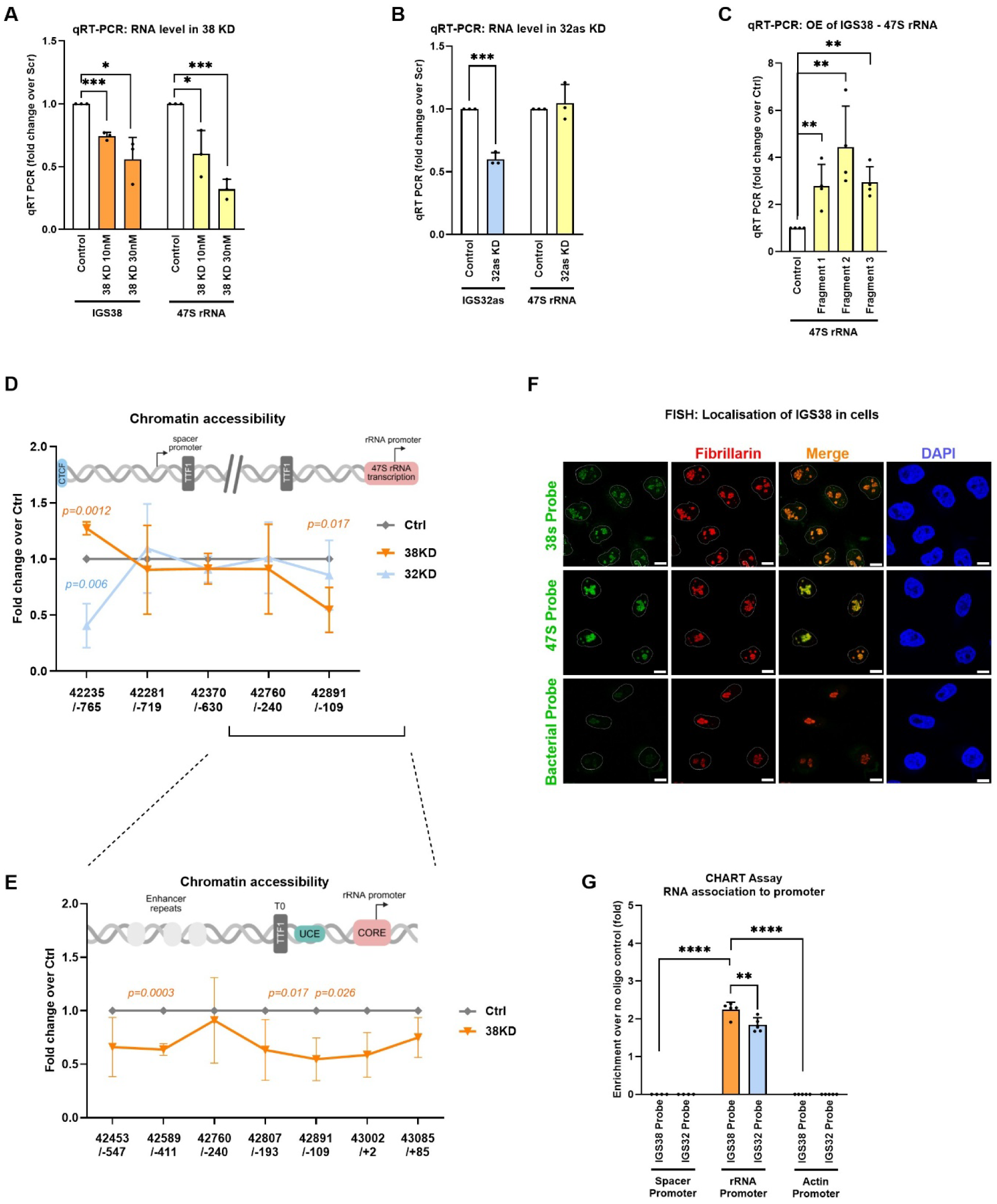
IGS38 regulates 47S gene expression and IGS38 knock down changes the chromatin compaction at the 47S rRNA gene promoter. A) Analysis of the 47S rRNA level after knock down of IGS38 using LNA-DNA Gapmers at 10 and 30 nM. cDNA from total RNA was normalised to 18S rRNA and the RNA levels are expressed relative to levels in cells treated with control LNA-DNA. B) Analysis of the 47S rRNA level after knock down of IGS32as. cDNA from total RNA was normalised to 18S rRNA. RNA levels are expressed relative to levels in cells treated with control LNA-DNA. C) Overexpression of IGS38 with three different fragments: Fragment 1, 200 bp from location 38.759 bp to 38.957 bp; Fragment 2, 200 bp from location 38.958 kb to 39.156 kb; Fragment 3, 400 bp from 38.756 bp to 39.156 bp, analysed for 47S rRNA levels by qRT-PCR. Normalisation was done to 18S rRNA and the RNA levels are related to the level in cells transfected with control vector. D) ATAC-qPCR analysis to detect chromatin accessibility at the spacer promoter, enhancer region and the 47S rRNA promoter in IGS38 knock down cells, IGS32as knock down cells or control cells. Signal values were normalised to the 27 kb region and presented as knock downs related to control. E) ATAC-qPCR data zoomed in at the enhancer region and 47S rRNA promoter region in IGS38 knock down cells or control cells. F) RNA-FISH localisation of IGS38 RNA in the nucleolus. Probes against 47S rRNA were used as a positive control and a bacterial probe as a negative control. Fibrillarin was used as a nucleolar marker and the merge shows fibrillarin and the RNAs (IGS38 or 47S rRNA) together, DAPI marked the nucleus. The scale bar marks 10 μm. G) CHART analysis using specific probes against IGS38 and IGS32as ncRNA at the spacer promoter and the rRNA promoter. The actin promoter was used as a negative control region. Data is presented as fold enrichment after normalising to no oligo control. All experiments present means for at least 3 biological replicates. Error bars represent the standard deviation. P-values are calculated with unpaired student’s t-test: * p≤ 0.05, **p≤ 0.01 and *** p≤ 0.001.

### IGS38 localises to the nucleolus and interacts with the rRNA gene promoter

Next, we determined the localisation of IGS38 by RNA-FISH with DIG labelled probes for IGS38 (sequences in Supplementary Table 1). The signal for the FISH probes selected for the IGS38 RNA co-localised with fibrillarin in the nucleolus in a manner similar to that of the 47S rRNA, even if less intense (Figure 2F). A bacterial probe was used as negative control (Figure 2F). We conclude that IGS38 remains in the nucleolus and regulates the 47S transcription. To assess whether the IGS38 RNA associated directly with nucleolar rRNA gene promoters, we designed oligonucleotide probes for chromatin hybridisation of RNA targets (CHART) assay of the IGS38 (nt from position 38.841 kb to 38.860 kb) and the IGS32as (nt from position 32.701 kb to 32.749 kb) (Sequences in Supplementary Table 1). The probes were assessed for specificity by depleting target ncRNAs using the RNase H enzyme (Supplementary Figures S2B), prior to determining the associations of the RNAs to the rRNA promoters and to their locations of origins in the IGS. The IGS38 associated with the 47S gene promoter but not with the spacer promoter downstream of the CTCF-boundary site (approximately -765 bp upstream of the TSS) (Figure 2G). The IGS32as also associated with the 47S rRNA promoter but not with the spacer promoter (Figure 2G). Furthermore, neither of these ncRNAs associated with the 38kb region, but both were found at the 32kb site (Supplementary Figure S2C). These results suggest that IGS38 moved away from its site of origin to the 47S rRNA gene promoter and to the 32kb region, in contrast to IGS32as which remained at its site of origin in 32 kb region. Taken together, IGS38 associated with the 47S rRNA gene promoter and affected the chromatin state to maintain an open configuration, whereas IGS32as had its main effect at its site of origin.

### IGS38 interacts with WSTF in the B-WICH complex at the 47S rRNA promoter

Since the association of IGS38 with the 47S rRNA promoter led to a more open chromatin at the region, we assessed its association with the WSTF protein and with the CSB protein, which both are chromatin remodelling proteins involved in activation of rRNA genes. The IGS38 interacted with WSTF-associated chromatin, but only to a lower level to CSB-associated chromatin (Figure 3A), whereas IGS32as associated to a lower level with both WSTF and CSB associated chromatin (Figure 3A). This indicates that IGS38 works with the B-WICH complex. To assess the direct binding, we next performed RNA IP and WSTF in particular displayed a high binding of IGS38, but not of IGS32as, while CSB bound IGS38 and IGS32as (Figure 3B).

**Figure 3.**
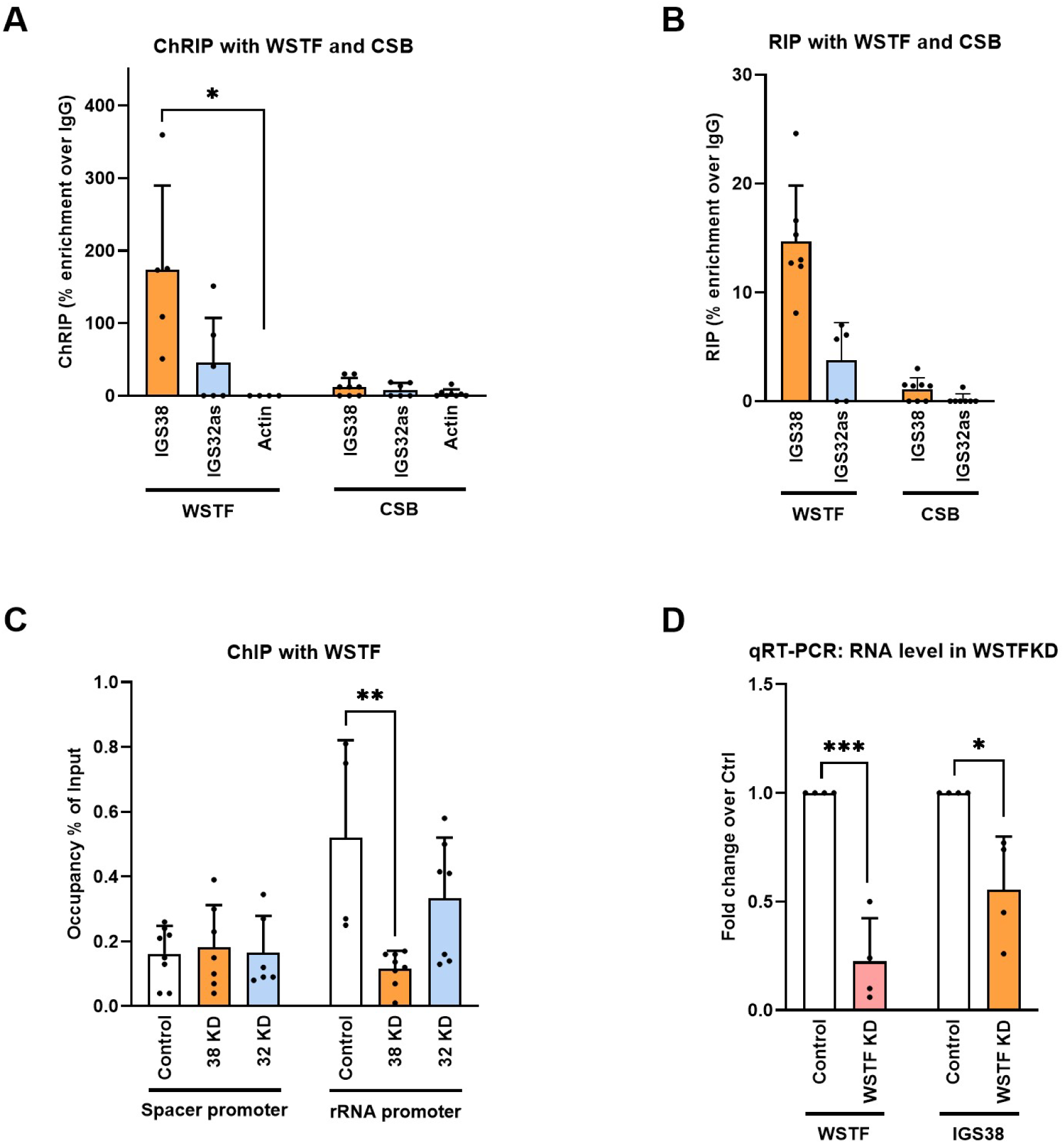
IGS38 recruits WSTF at the 47S rRNA promoter. A) ChRIP analysis of IGS38 and IGS32as association with B-WICH bound chromatin using antibodies against WSTF and CSB. Actin was used as a negative control. qRT-PCR data is shown as percentage enrichment over IgG. B) RNA IP to analyse direct interactions between IGS38 and IGS32as with WSTF and CSB. qRT-PCR data is shown as percentage enrichment over IgG. C) ChIP-qPCR of WSTF occupancy at the spacer promoter and 47S rRNA promoter under IGS38 knock down cells, IGS32as knock down cells or control cells. Signals are calculated as percentage of input samples. D) qRT-PCR of WSTF and IGS38 levels in WSTF knock down cells related to control cells. The RNA is normalised to 18S rRNA. All experiments present means for at least 3 biological replicates. Error bars represent standard deviation. P-values are calculated with unpaired student’s t-test: * p≤ 0.05, **p≤ 0.01 and *** p≤ 0.001.

Next, we performed structural analyses assessing minimum free energy (MFE) algorithms for base pair probabilities and positional entropy of the sequences between 38.8 kb and 39.1 kb of IGS38 (Supplementary Figure S3A). This region of the IGS38 formed stretches of hair-pin structures and stabilised bulges with high confidence, predicted to interact with proteins or RNA (63, 64). WSTF, in contrast to CSB, is an RNA binding protein and binds the ETS region of the 47S transcript (39, 30), which led us to use the catRAPID omics v2.0 (65) to assess specific protein-RNA interactions between the WSTF and IGS38. The WSTF was predicted to bind at a large region of the RNA from base pair 120 to 200, corresponding to sequences from approximately position 38.9 to 39.0 kb in the IG3S38 target sequence (Supplementary Figures S3B), found in the long hairpin loop pointing downwards in the predicted structures (Supplementary Figures S3A). IGS32as was only predicted to bind at a narrow stretch at the one end of the RNA and IGS19as did not bind WSTF at all (Supplementary Figures S3C to S3D). The binding of the IGS38, and to a certain extent IGS32as, mapped to regions in the whole WSTF protein; with a higher propensity score at a region around amino acid 750, which is between the DDT domain and the PHD domain of the WSTF where a few internally disordered domain regions reside (https://www.ebi.ac.uk/interpro/protein/UniProt/Q9UIG0/) as well as the BAZ1, BAZ2 and the WACZ domains (66). Taken together, we conclude that IGS38 interacts directly with WSTF. We also investigated the Treacle protein (encoded from the TCOF gene), which is involved in activating rRNA gene expression (67), but this protein did not bind to chromatin-associated IGS38 or IGS32as, or directly to the RNA (Supplementary S3E and S3F), which is supported by the low predicted binding levels obtained from CatRAPID (Supplementary Figure S3G). We conclude that IGS38 binds to WSTF through specific domains and suggests that this interaction is important in recruiting the B-WICH complex to the 47S promoter to alter the chromatin configuration.

### IGS38 and WSTF associate at the 47S rRNA promoter

To assess the functional interaction between WSTF and IGS38 at the promoter, we knocked down the IGS38 RNA and detected a reduced association of WSTF at the 47S rRNA promoter but not at the spacer promoter (Figure 3C). We also investigated whether B-WICH regulated the IGS38 transcript, since the B-WICH complex component WSTF and SNF2h associated with the 38 kb region (Supplementary Figure S3H). Indeed, WSTF regulated the transcription of IGS38, as WSTF knock down reduced the level of IGS38 by 60% (Figure 3D). This is also reflected in the fragments generated from the 37 kb region, which were reduced by WSTF knock down in HeLa cells (Supplementary Figure S3I). This indicates a mechanistic control of the 47S rRNA gene transcription by a WSTF – IGS38 regulation loop.

### IGS38 knock down reduces the level of UBF at the 47S rRNA promoter

The changed chromatin accessibility at the 47S rRNA gene caused by IGS38 reduction prompted us to investigate whether these changes were associated with alteration in the chromatin landscape by ChIP. IGS38 knock down resulted in a significant reduction in UBF at the 47S rRNA promoter, but not at the spacer promoter, whereas IGS32as knock down had no effect on the UBF level at any of the promoters (Figure 4A). We did not detect any change in histone H3 levels upon IGS38 or IGS32as knock down at the promoters (Figure 4B), while histone H2A.Z, mainly observed in active chromatin, increased at the spacer promoter in IGS38 knock down (Figure 4C). The association of CTCF marks a border element between the chromatin at the promoter-enhancer region and the chromatin in the IGS (5, 47, 68). The level of CTCF was reduced at the border element at the spacer promoter in IGS38 knock down cells and increased at the 47S rRNA promoter (Figure 4D), indicating dysregulation of the chromatin in particular at the spacer promoter. TTF1, which blocks read-through transcripts (13) and is involved in the RNA:DNA hybrids generated between silencing RNAs and DNA at the promoter, was also changed at the region in IGS38 knock down cells; it was reduced at the 47S rRNA promoter in IGS38 knock down, but not at the Tsp site downstream of the spacer promoter (Figure 4E). The presence of TTF1 at the spacer promoter indicates that it still blocked read-through transcripts from the spacer promoter, as the TTF1 does at the Tsp in mouse cells (13).

**Figure 4.**
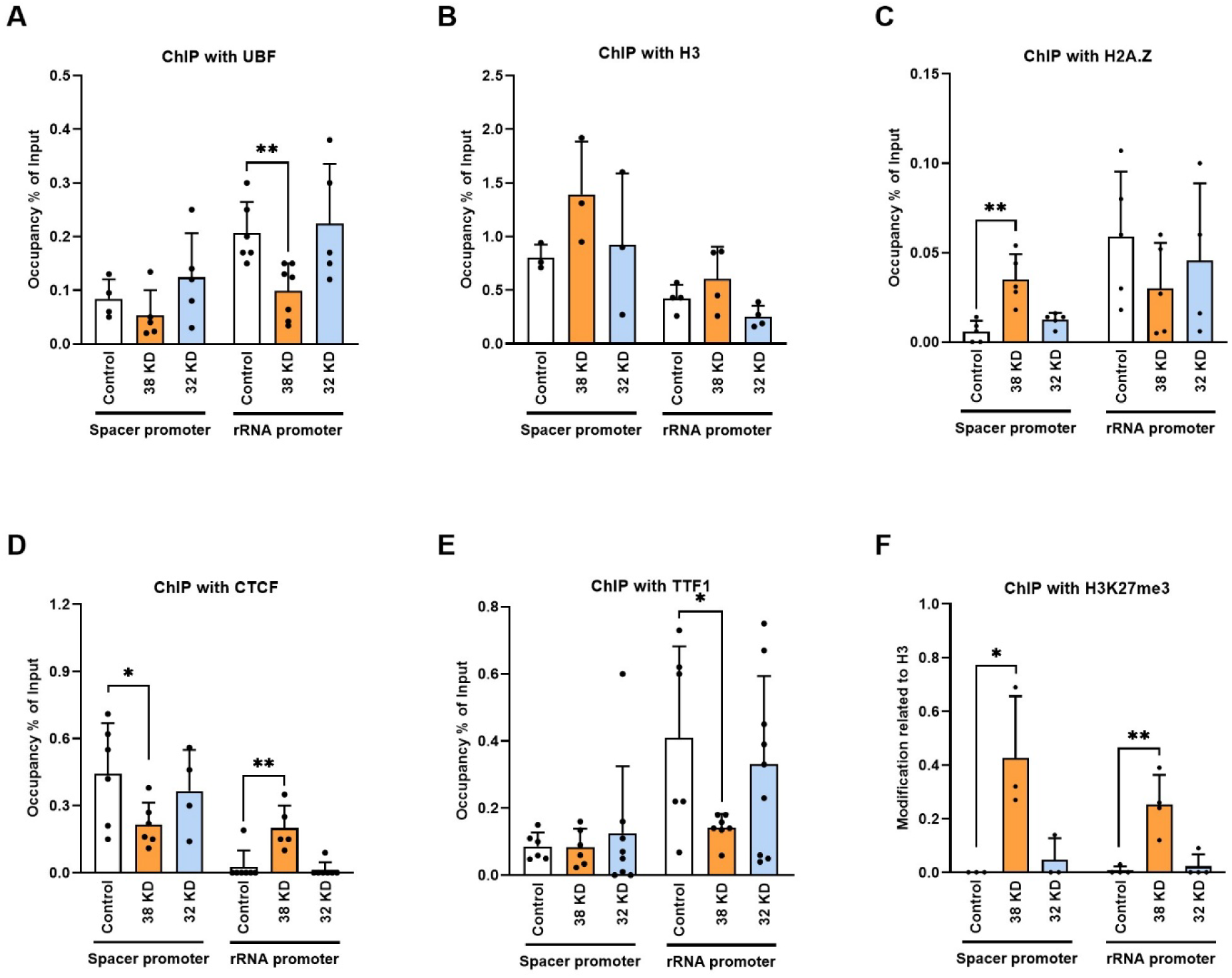
Chromatin factors at the spacer and 47S rRNA gene promoter in IGS38 and IGS32as knock down. A) ChIP-qPCR analysis of UBF occupancy at the spacer promoter and rRNA promoter in control cells, IGS38 knock down cells, and IGS32as knock down cells. B) ChIP-qPCR analysis of histone H3 occupancy at the spacer promoter and rRNA promoter in control cells, IGS38 knock down cells, and IGS32as knock down cells. C) ChIP-qPCR analysis of histone H2A.Z occupancy at the spacer promoter and rRNA promoter in control cells, IGS38 knock down cells, and IGS32as knock down cells. D) ChIP-qPCR analysis of CTCF occupancy at the spacer promoter and rRNA promoter in control cells, IGS38 knock down cells, and IGS32as knock down cellsE) ChIP-qPCR analysis of TTF1 occupancy at the spacer promoter and rRNA promoter in control cells, IGS38 knock down cells, and IGS32as knock down cells. All signals from A) - E) are calculated as percentage of input samples. F) ChIP-qPCR analysis of the presence of the histone H3K27me3 at the spacer promoter and rRNA promoter in control cells, IGS38 knock down cells, and IGS32as knock down cells. The signal is relative to the histone H3 signal. All experiments present means for at least 3 biological replicates. Error bars represent the standard deviation. P-values are calculated with unpaired student’s t-test: * p≤ 0.05, **p≤ 0.01 and *** p≤ 0.001.

Furthermore, H3K27me3 modified histone H3 was significantly higher in IGS38 knock down cells at both the spacer and the 47S rRNA gene promoter (Figure 4F). H3K27me3 is established by EZH2 in the PRC2 complex and is often accompanied with PRC1 to induce transcriptional repression. The PRC1 complex deposits H2AK119ub which stabilises PRC2 and the conventional PRC1 component CXB2 (69, 70). Despite the increase of nucleosomal H3K27me3 in IGS38 knock down cells, no increase in PRC1-associated factors was observed; rather a decrease of nucleosomal H2AK119ub at the 47S rRNA promoter with no change of the canonical PRC1 protein CBX2 was observed (Supplementary Figures S4A and S4B). H3K27me3 and Polycomb-silencing is rarely found at rDNA and the dysregulated H3K27me3-level suggests that a Polycomb-independent EZH2 activity was operating in IGS38 knock down cells. Taken together, a reduced IGS38 level disrupts the chromatin organisation at the promoters which results in a loss of UBF at the 47S rRNA gene promoter.

### IGS38 modulates PIC formation at the 47S rRNA gene promoter

Next, we investigated whether IGS38 knock down affected the binding of the rRNA transcription machinery and the formation of the PIC (polymerase initiation complex) at the promoters. Using primer pairs covering to the transcribed rRNA gene copy (depicted in Figure 5A), we detected an accumulation of RNA pol I at the 47S rRNA gene promoter in IGS38 knock down cells compared to control cells (Figure 5B) while the occupancy at the ETS region (1 kb downstream of the TSS) was lower. The level was similar further into the gene body, 4 kb (18S) and 8 kb (28S) from the TSS, and Transcription Termination site (TTE) at the end of the genes (13 kb) (Figure 5C). IGS32as knock down did not lead to an accumulation of RNA pol I at the promoter, only to a lower occupancy of at the ETS (Figure 5B and 5C). The accumulation of RNA pol I at the promoter with a lower occupancy in the ETS region in IGS38 knock down cells suggests that a large fraction of RNA pol I was stalled at the promoter. Generally, RRN3/TIF1A interacts with the RNA pol I to allow the formation of a transcriptionally competent PIC and to increase its recruitment to the promoter (71–73). In fact, we found a significant decrease in RRN3 recruitment in IGS38 knock down cells to the 47S rRNA promoter (Figure 5D), suggesting that the stalled RNA pol I was transcriptionally incompetent. These changes in RNA pol I and RRN3 were not detected at the spacer promoter (Figures 5B and 5D). Next, we investigated the occupancy of SL1 components, TBP and TAF1C, which did not change significantly at any of the promoters in IGS38 or IGS32as cells (Figure 5E and 5F). Taken together, we conclude that IGS38 facilitates the interaction of RRN3 with RNA pol I and promotes the formation of a transcription competent RNA pol I and promoter escape.

**Figure 5.**
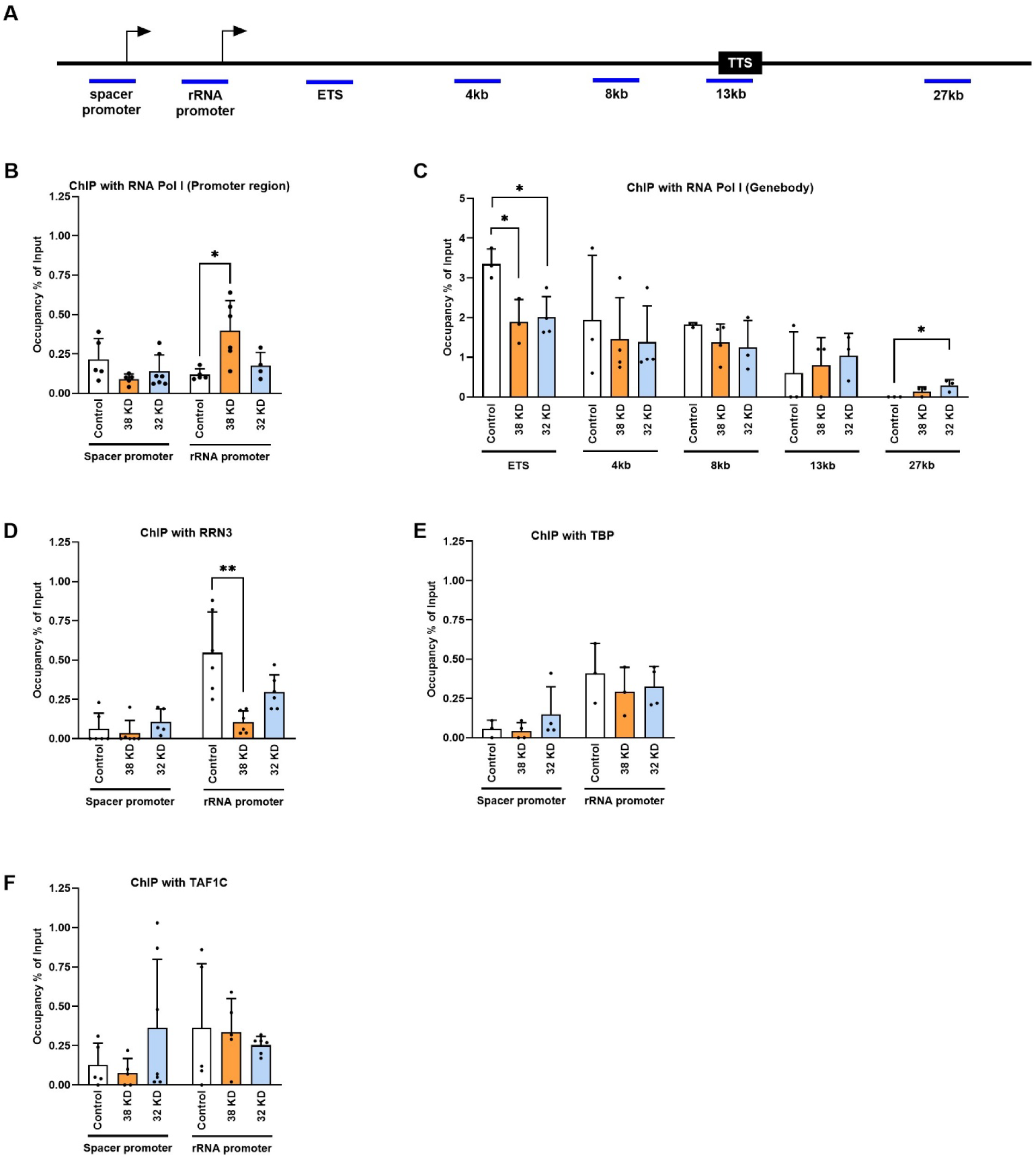
Recruitment of PIC components at the 47S rRNA gene promoter. A) Graphical scheme of the locations of the primers used over the rDNA gene repeat: spacer promoter, rRNA promoter, ETS (1 kb), 4 kb (18S), 8 kb (28S), 13 kb (Transcription terminator site, TTS) and 27 kb (in the IGS). B) ChIP-qPCR analysis of RNA pol I occupancy at the spacer promoter and 47S rRNA gene promoter in control cells, IGS38 knock down cells, and IGS32as knock down cells. C) ChIP-qPCR analysis of RNA pol I occupancy in the gene body with primers indicated in (A) in control cells, IGS38 knock down cells and IGS32as knock down cells. D) ChIP-qPCR analysis of RRN3 occupancy at the spacer promoter and 47S rRNA gene promoter in control cells, IGS38 knock down cells, and IGS32as knock down cells. E) ChIP-qPCR analysis of TBP occupancy at the spacer promoter and 47S rRNA gene promoter in control cells, IGS38 knock down cells, and IGS32as knock down cells. F) ChIP-qPCR analysis of the SL1 component TAF1C occupancy at the spacer promoter and 47S rRNA gene promoter in control cells, IGS38 knock down cells, and IGS32as knock down cells. The signals in all panels are calculated as percentage of input samples. All experiments present means for at least 3 biological replicates. Error bars represent the standard deviation. P-values are calculated with unpaired student’s t-test: * p≤ 0.05 and **p≤ 0.01.

### IGS38 binds to the SL1 an RRN3 at the promoter

Given the association of IGS38 with chromatin harbouring the RNA pol I activation factors UBF and WSTF (Figure 1F and 3A, respectively), we investigated the interaction with further RNA Pol I factors. IGS38 but not IGS32as interacted with TAF1C and RRN3 in chromatin (Figure 6A), and given that these factors are binding at promoters we conclude that IGS38 associates with the RNA pol I factors at the RNA promoters. The poor sequence homology between the rRNA promoter (200 bp upstream of the TSS and the 400 bp of the core of IGS38) displaying only 30% identity (Supplementary Figure S6A) argues against a mechanism of forming a direct RNA:DNA hybrid, and led us instead to investigate direct binding to factors at the promoters. IGS38 bound directly to TAF1C and RRN3, but not to UBF, whereas IGS32as did not bind to any of the factors (Figure 6B). The cancer specific LINC01116, originating from its locus on chromosome 2 in the nucleoplasm, also interacts with SL1 and enhances transcription (45). Therefore, we investigated the binding of LINC01116 and SLERT, an activating DDX21 binding lncRNA (46), to RNA Pol I factors. LINC01116 bound to TAFIC, RRN3 and WSTF, whereas SLERT only displayed a strong binding to DDX21 (Supplementary Figure S6B). The interaction between LINC01116 and WSTF was also predicted by a catRAPID analysis, but with a narrower pattern to that of IGS38 and only to a small region of the RNA (Supplementary Figure S6C). SLERT was not predicted to bind to WSTF at all (Supplementary Figure S6D). We propose that IGS38, but not IGS32as, interacts with RNA pol I factors SL1 and RRN3 to stabilise PIC formation and with WSTF in the B-WICH complex to open the chromatin at the 47S rRNA promoter, which leads to an enhanced 47S rRNA gene transcription in growing cells (Figure 6C). This mechanism may be replicated by certain cancer associated RNA species, as LINC01116 also binds to SL1 and WSTF to enhance 47S rRNA transcription.

**Figure 6.**
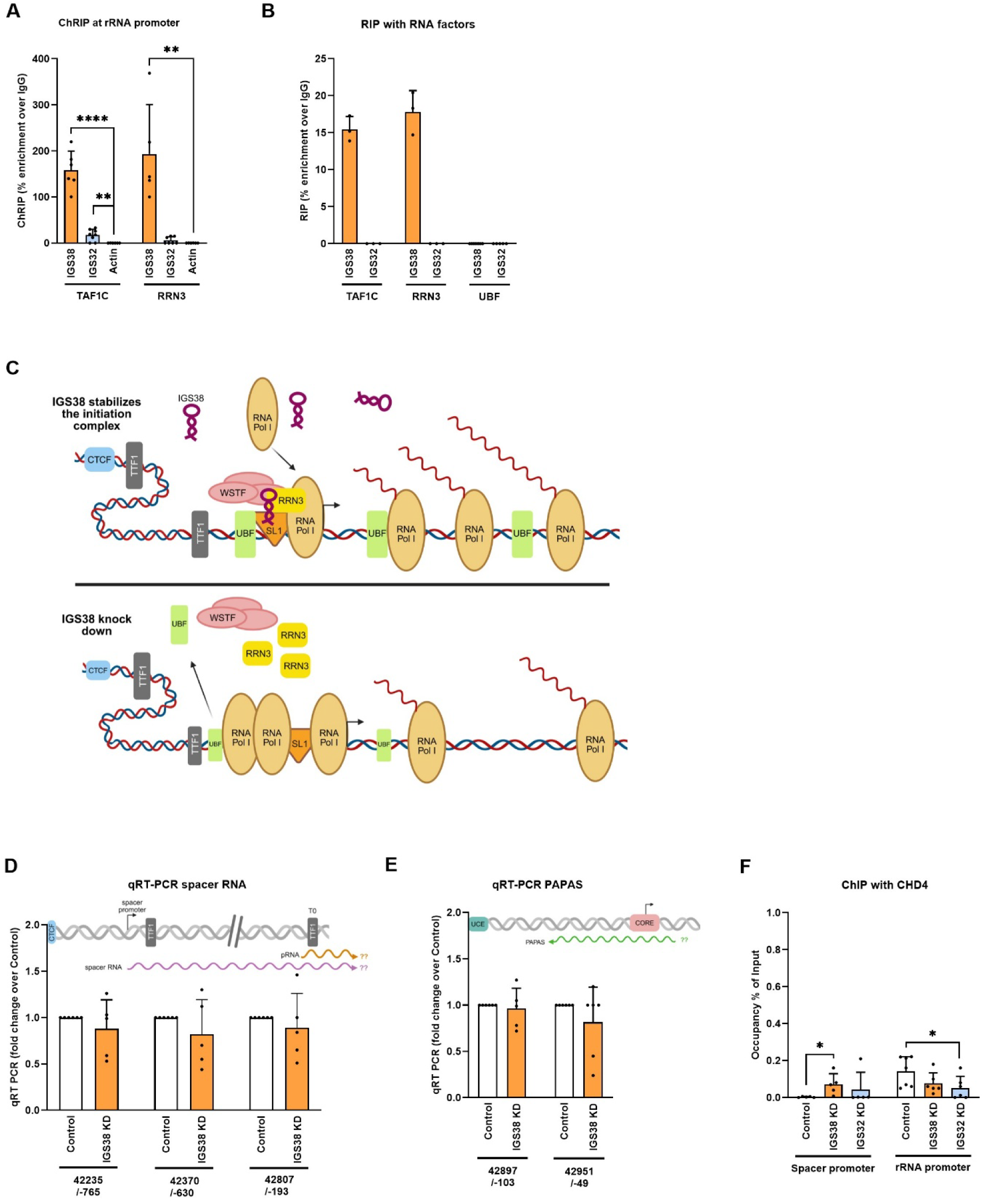
IGS38 interacts with the promoter factors SL1 and RRN3 at the promoter. **A)** ChRIP of the association of IGS38 and IGS32as with TAF1C and RRN3 in chromatin. Actin was used as a negative control. RT-qPCR data is shown as percentage enrichment over IgG. B) RNA IP to analyse direct interactions between IGS38 and IGS32as with TAF1C, RRN3 and UBF. RT-qPCR data is shown as percentage enrichment over IgG. C) Model of the action of IGS38 and WSTF at the 47S rRNA gene promoter in growing cells (top) compared to IGS knock down cells (bottom). In growing cells IGS38 stabilises RNA Pol I in a transcription competent state, whereas in IGS38 knock down cells UBF levels are reduced, RRN3 is lost together with WSTF and concurrent the RNA pol I is accumulating at the promoter. Created by biorender (https://biorender.com/shortURL). D) Spacer RNA transcript expression at the spacer promoter (location 42235/-765), downstream of the TSS of the spacer-RNA (location 42370/-630), and at the T0 site (location 42807/-193), binding site of the pRNA. The qRT-PCR data was normalised to 18S rRNA and shown as fold change over Gapmer control. E) qRT-PCR of IGS38 KD and control cells to determine RNA levels of the of PAPAS expression at the position 42897/-103 at the 47S RNA promoter and 42951/-49 over the TSS of the 47S rRNA. The data was normalised to 18S rRNA and shown as fold change over Gapmer control. F) ChIP-qPCR analysis of CHD4 occupancy at the spacer promoter and 47S rRNA gene promoter in control cells, IGS38 knock down cells, and IGS32as knock down cells. The signals are calculated as percentage of input samples. All experiments present means for at least 3 biological replicates. Error bars represent the standard deviation. P-values are calculated with unpaired student’s t-test: * p≤ 0.05, ** p≤ 0.01, *** p≤ 0.001 and **** p≤ 0.0001.

### IGS38 works independently from silencing ncRNA spacer-RNA and PAPAS

The spacer-RNA from the spacer promoter and the antisense PAPAS from the 3’ end of the transcribed gene both establish silent chromatin states (16, 21). Since the IGS38 increases the 47S rRNA transcription, we asked whether IGS38 acted by a mechanism that counteracted these silencing rRNAs. However, we did not detect any changes in the already low levels of spacer-RNA transcripts at the enhancer region upon IGS38 knock down (Figure 6D), not even at the designated pRNA region which transcript binds to the T0 (16, 21). We conclude that IGS38 is not involved in the control of the transcription of the spacer-RNA. Antisense PAPAS read-through transcripts at the 47S rRNA gene promoter region is high in cells in which 47S gene transcription is low, as in different stress conditions or differentiation (35). Despite a reduced 47S rRNA gene transcription concomitant with a closed chromatin state at the promoter in IGS38 knock down cells, no higher level of PAPAS upstream of the TSS was observed (Figure 6E). Furthermore, PAPAS transcripts recruit CHD4 or Suv4-20h2 to the promoter during different stress conditions, either by forming triple helices or producing R-loops at the promoter (21, 33, 35). We therefore investigated whether IGS38 knock down led to higher level of R-loops by DRIP (DNA-RNA immunoprecipitation) with the antibody S9.6. We did not detect any R-loops in the gene or at the site of origin of our identified ncRNAs unless we challenged the transcription by 3 hours glucose stimulation after glucose starvation (Supplementary Figure S6E). R-loops were formed upon stimulation at 19 kb, 32 kb and in the 28S part of the gene, but not at the 38 kb site. Nevertheless, we did not detect any R-loops at the promoters or at the end of the transcribed gene in IGS38 or IGS32as knock down cells (Supplementary Figure S6F). Furthermore, we only detected low occupancy of CHD4 or Suv4-20h2 at the promoters in control cells and no difference was found in IGS38 and IGS32as knock down cells, except for a small, but significant, increase of CHD4 at the spacer promoter in IGS38 knock down cells (Figure 6F and Supplementary Figure S6G). Taken together, there is no interplay between the silencing RNAs species and IGS38 in growing cells and the transcription of the PAPAS or the spacer-RNA do not only depend on reduced 47S rRNA transcription.

### Cellular stresses differentially modulate IGS transcripts

PAPAS is formed in response to various stressors and recruits CHD4 in the NuRD in response to heat shock and hypotonic stress (22, 24, 26) and Suv4-20h2 in response to growth factor depletion (25). As we did not observe any correlation between the functions of IGS38 and PAPAS in growing cells, we examined the response of the lncRNAs to stress conditions. IGS38 only responded to hypotonic stress, which resulted in a reduction of the transcript (Figure 7A), while no change in IGS38 was observed during growth factor depletion or heat shock (Supplementary Figures S7A and S7B). The levels of PAPAS displayed a trend to an increase in all conditions, with a significant induction, 7-fold, upon hypotonic stress (Figure 7A and Supplementary Figures S7A and S7B). IGS32as did not change in any conditions, while IGS19as was induced by growth factor depletion and heat shock, and less upon hypotonic stress (Figure 7A and Supplementary Figure S7A and S7B). The association of WSTF and the binding of RNA pol I to the 47S rRNA promoter was also severely reduced, with only a weak increase in the recruitment of CHD4, upon hypotonic stress compared to control cells (Figure 7B). The association of WSTF with the promoter was also reduced in growth factor depleted cells, but with an increase of Suv-20h2 (Supplementary Figure S7C). The difference in response of IGS38 and WSTF to the stressors suggests that they also work independently.

**Figure 7.**
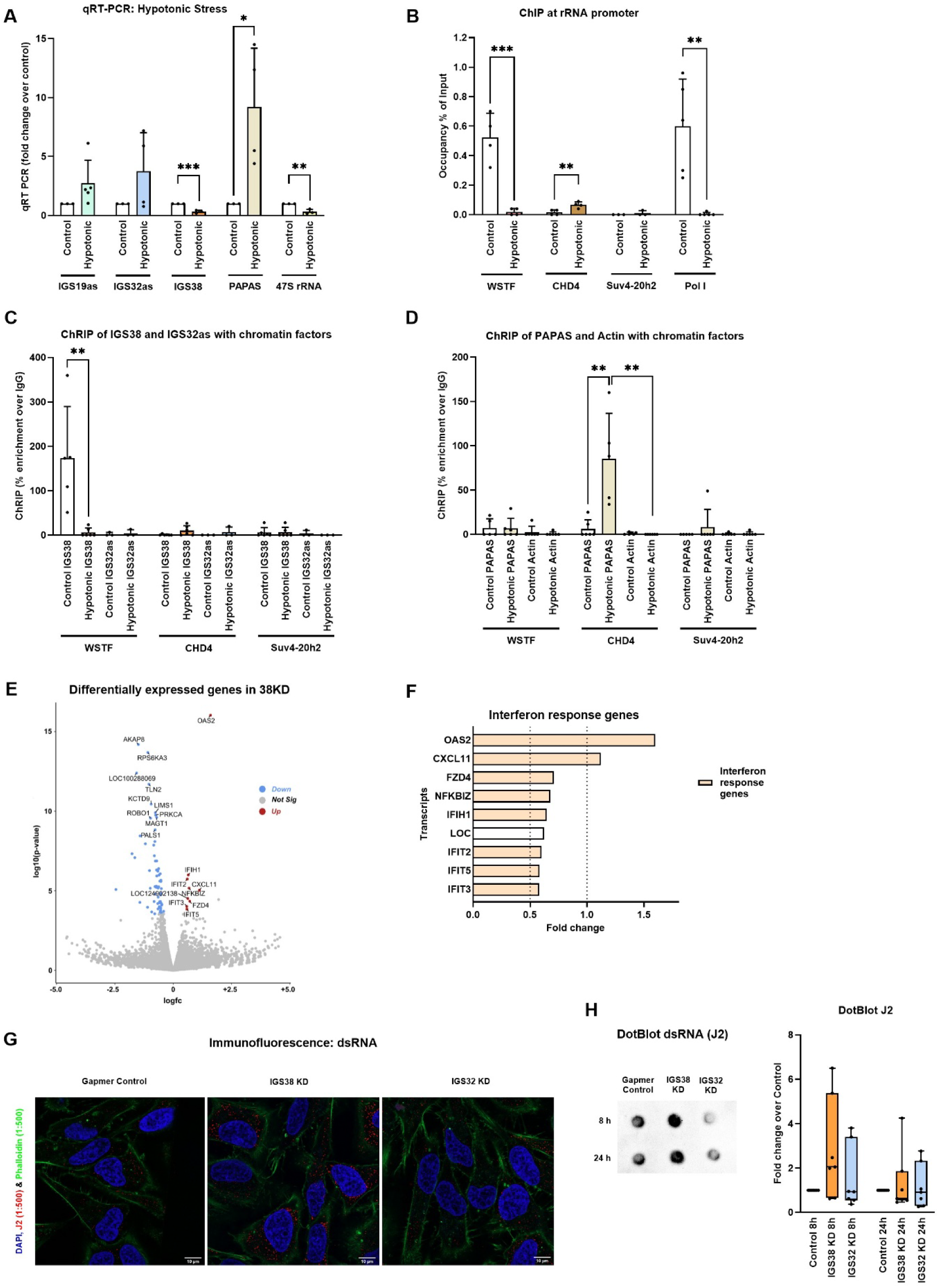
Noncoding RNAs from human rDNA respond differently to stress and IGS38 KD leads to accumulation of dsRNA in the cells. A) qRT-PCR of the RNA levels of IGS19as, IGS32as, IGS38, PAPAS and 47S rRNA in cells exposed to hypotonic stressed and control cells. The data was normalised to 18S rRNA and shown as fold change over control. B) ChIP-qPCR analysis of WSTF, CHD4, Suv4-20h2 and Pol I occupancies at the 47S rRNA gene promoter in cells exposed to hypotonic and control cells. The signals are calculated as percentage of input samples. C) ChRIP of the association IGS38 and IGS32as with chromatin bound WSTF, CHD4 and Suv4-20h2 in cells exposed to hypotonic stress and control cells. RT-qPCR data is shown as percentage enrichment over IgG. D) ChRIP of the association of PAPAS with chromatin bound WSTF, CHD4 and Suv4-20h2 in cells exposed to hypotonic stress and control cells. The actin RNA was used as a control. RT-qPCR data is shown as percentage enrichment over IgG. E) Volcano plot for RNA sequencing of IGS38 knock down cells to control showing significant differentially expressed genes. F) Genes in the Gene Ontology term Interferon response genes, differentially expressed in IGS38 knock down cells compared to Gapmer control cells. **G)** Immunofluorescence in control and IGS38 knock down cells stained with J2-antibody against dsRNA (red), FITC-phallodin actin filaments in the cytoplasm (green), and DAPI staining the nucleus (blue). The size bars in the images represent 10 μm. **H)** The abundance of dsRNA by Dot Blot in Gapmer control cells, IGS38 knock down cells, and IGS32as knock down cells using the J2 antibody at 8 hours and 24 hours as indicated above the image in the left panel. Quantification of the dsRNA levels in the samples are shown at 8 hours and 24 hours are shown in the right panel. The median value in each box is marked. All experiments present means for at least 3 biological replicates. Error bars represent the standard deviation. P-values are calculated with unpaired student’s t-test: * p≤ 0.05, ** p≤ 0.01 and *** p≤ 0.001.

### IGS38 does not bind silencing factors CHD4 or Suv4-20h2

To further examine the underlying mechanism of IGS38, we investigated which factors associated with the lncRNAs upon hypotonic stress. The interaction of WSTF with chromatin-associated IGS38 was lost upon hypotonic stress, which may reflect the low abundance of IGS38 (Figure 7C). CHD4 and Suv4-20h2 did not interact with chromatin-associated IGS38 (Figure 7C). In addition, the factors TAF1C, RRN3, and UBF were lost, all of which were associated with chromatin-associated IGS38 in growing cells (Supplementary Figure S7D). TAF1C and RRN3 did not associate with chromatin associated IGS32as in growing cells, but UBF was lost during hypotonic stress. (Supplementary Figure S7D). Since the PAPAS level was higher upon hypotonic stress, we also investigated the association of factors to chromatin-associated PAPAS. Similar to growing cells, WSTF did not associate to the chromatin-associated PAPAS during hypotonic stressed (Figure 7D). In line with the finding that CHD4 is recruited to the 47S rRNA promoter by PAPAS, CHD4 was enriched on chromatin-associated PAPAS in hypotonic stress, while Suv4-20h2 did neither associate in growing cells nor in cells during hypotonic (Figure 7D). The RNA pol I factors TAF1C, RRN3, and UBF interacted at low levels with chromatin-associated PAPAS in growing cells and these factors remained low upon hypotonic stress (Supplementary Figure S7E). Furthermore, despite that CHD4 was interacting with chromatin-associated PAPAS, we could not detect any direct binding with PAPAS (Supplementary Figure S7F). Neither did any of the other factors, WSTF, Suv4-20h2, UBF, TAF1C or RRN3, bind PAPAS (Supplementary Figure S7F). Additionally, no CatRAPID predicted binding of PAPAS to WSTF was detected as the propensity value was negative (Supplementary Figure S7G). We conclude that PAPAS transcripts and the IGS38 RNA do not function together and are differentially regulated during stress.

### IGS38 is expressed at the same levels in primary cells and cancer cell lines

The abundance of PAPAS is higher in differentiated cells than in proliferating cells and cancer cells (35), which prompted us to examine the abundance of IGS38 and PAPAS in different cancer cells lines and in primary immune cells and fibroblast. The abundance of PAPAS was lower than that of IGS38; approximately 1-2% of the level of IGS38 in HeLa cells. HeLa cells and human osteosarcoma U2OS cells expressed approximately the same levels of the two RNAs whereas human mammary epithelial MCF10A cells and the embryonic kidney HEK293 cells expressed lower levels of IGS38. MCF10A cells expressed higher levels of PAPAS than the other cells, and embryonic kidney HEK293 cells expressed very low levels of PAPAS (Supplementary Figure S7H). The primary cells investigated, fibroblasts, monocytes and monocyte derived dendritic cells (moDCs), expressed approximately the same levels of IGS38 as the cell lines, but had higher levels of PAPAS (Supplementary Figure S7I). High growth rate and expression of PAPAS are correlated with a higher expression of RPA (replication protein A) and RNAse H1, which resolve R-loops and counteract PAPAS formation, for instance in cancer cells (34, 35). The RPA2 isoform and RNAse H1 had the same abundance in HeLa and U2OS, but only the RPA2 was induced in both cell lines upon hypotonic stress (Supplementary Figure S1J). This increase of RPA2 may be part of a stress response to counteract the sudden silencing of rRNA genes by PAPAS and NuRD in cancer cells.

### Reduction in IGS38s levels is associated with a weak dsRNA Interferon response

Most intergenic transcripts from the rDNA loci have a function in nucleolar transcriptional regulation or cellular stress response (39, 40) but may have other functions. To determine whether IGS38 RNA affected the expression of other genes, we performed an RNA sequencing analysis of IGS38 knock down in HeLa cells. Comparing the knock down with control cells showed that only a small number of transcripts was significantly differentially expressed in IGS38 knock down cells (Figure 7E). Gene ontology analysis for biological processes revealed that most of the transcripts that show higher expression in the IGS38 knock down samples were related to antiviral innate immune response (GO:0140374), negative regulation of viral genome (GO:0045071), and cytokine mediated signalling pathway (GO:0019221), with OAS2 (2’-5’ oligoadenylate synthetase) and CXCL11 as the most significantly upregulated RNAs (Figure 7F). We validated the increased expression of OAS2 in IGS38 knocked down in HeLa cells by qRT-PCR, and an increase also occurred in U2OS cells showing that this effect was not limited to HeLa cells (Supplementary Figure S7K). The OAS enzymes are dsRNA sensors which activate RNase L in response to dsRNA in the cytoplasm (74, 75), and interestingly, a further cytoplasmic dsRNA sensor, MDA5 (IFIH1 gene), was also slightly upregulated together with interferon stimulated genes (ISGs) in response to a reduced level of IGS38. This prompted us to investigate whether IGS38 knock down resulted in an increase of dsRNA abundance in the cytoplasm by performing immunofluorescence using the J2-anti-dsRNA antibody in cells transfected with LNA-control, LNA-IGS38 and LNA-IGS32as. IGS38 knock down resulted in a higher abundance of dsRNA accumulating in the cytoplasm than in IGS32as knock down cells or control cells (Figure 7G). We next assessed the abundance of dsRNA in these cells by isolating dsRNA at different time points and knock down of IGS38, but not IGS32as to the same extent, resulted in higher abundance of dsRNAs, as early as 8 hours (Figure 7H). However, the higher dsRNA level in IGS38 cells was not sufficient to activate RNAse L, which degrades all RNA when activated by OAS enzymes (Supplementary Figure S7L). Nevertheless, this suggests that dysregulation of IGS38 leads to higher dsRNAs in the cytoplasm which could potentially stimulate a cellular immune response.

## Discussion

We have identified three lncRNAs originating from the IGS in the rDNA loci of human cells. These lncRNA could be isolated as RNA species longer than 500bp and have different functions. In addition to a stress related lncRNA, IGS19as, from the region downstream of the gene termination site, two additional lncRNAs, the chromatin-associated IGS38 and IGS32as, were identified. Most of the previously reported nucleolar lncRNA are involved in stress responses, sequestering stress response protein (39, 40), or silencing of the rRNA gene expression, such as the spacer-RNA and PAPAS which recruit chromatin remodelling factors NoRC, NuRD and Suv4-20h2 (11–26). We show that the IGS38 ncRNA is involved in rRNA transcription; it associated with the rRNA gene promoter and in contrast to the lncRNAs spacer-RNA and PAPAS, it positively regulated 47S rRNA gene transcription. IGS38 activated transcription by acting as chromatin-associated scaffolding RNA to stabilise the RNA pol I machinery and by cooperating with the WSTF in the chromatin remodelling complex B-WICH. IGS38 bound to the RNA pol I factors SL1 and RRN3 directly at the promoter, but also to the WSTF protein to establish a more accessible 47S rRNA promoter. We propose that IGS38 promotes enhanced rRNA transcription, since IGS38 down regulation led to stalling of RNA pol I in a transcription incompetent state without RRN3 at the promoter, suggesting impaired promoter escape. Dysregulation of the IGS38 function led weak immune response to dsRNAs. The chromatin-associated RNA pol I IGS32as ncRNA had no effect on rRNA gene transcription, nor did it change in stress, instead it interacted with histone H3K9me3 and the heterochromatin protein HP1α. We propose that IGS32as is involved in the in maintenance of constative heterochromatin by a mechanism where RNA transcription is required for the recruitment of heterochromatin protein HP1α.

The IGS region in the rDNA is not conserved between species, but a comparison between primates displays a similar structure of functional region. The regions between 15 kb to 21 kb downstream of the TSS and between 28 kb and 33 kb downstream of the TSS, as well as the upstream region of the promoter are conserved between primates, and the region 37 kb to 40 kb also harbours elements of conservation, some small elements also with the mouse IGS (76). Transcripts have been identified from these regions and both RNA pol I and RNA pol II are present in the human rDNA repeats. In particular, RNA polymerases are present at the region 16 kb to 32 kb, from which stress response transcripts are generated, such as sincRNAs and asincRNAs (39, 40). The RNA pol II antisense IGS19as transcript originated from this region and is upregulated by different stressors, suggesting that it is most likely an asincRNA. The RNA pol II transcript IGS38 originated from the region around 37 kb to 39.5 kb. The initiating RNA pol II with serine 5 phosphorylated CTD (Ser5-pol II) and the elongating RNA pol II with serine 2 phosphorylated CTD (Ser2-pol II) as well as the kinase CDK9 have been found in two peaks in the IGS (40); in a region around 28 kb to 30 kb and in the 38 kb region, which transcription is important to form RNA:DNA hybrids to protect nucleolar integrity and rRNA biogenesis. We detected both Ser-5-RNA pol II and Ser2-RNA pol II in the 38.5 kb region, but no R-loops which indicates active transcription in this region of ncRNAs with other functions than nucleolar integrety. The IGS38 transcript was identified as a 1300 bp RNA, but it is likely that smaller fragments exist, either by degradation or by transcription from multiple promoters. We have identified two putative RNA pol II promoters in the 38 kb to 39 kb region: 38.5 and 38.7 kb from the TSS of the rRNA gene. Nevertheless, other promoter elements may be used by RNA pol II in the IGS; in the 28 kb region of the rDNA, RNA pol II uses alternative general RNA pol II factors, TBPL and PAF1, for transcription initiation and elongation (77). TBPL is involved in initiation of paused RNA pol II from TATA-less promoters, and PAF1 involved in pause release. These factors may also be involved in the transcription of IGS38 from alternative promoters.

The IGS38 was involved in the activation of the 47S rRNA transcription, but in contrast to the silencing spacer-RNA or PAPAS transcribed from the loci, did not act at the origin of transcription. The IGS38 transcript originated from a location around 38.9 kb from the TSS and associated with the 47S rRNA promoter, maintained the level of UBF and allowed for RRN3 to activate RNA pol I. UBF and SL1 have been proposed to co-operatively interact with each other to establish the right conformation at the promoters for RNA pol I binding (68, 71, 72, 72, 78, 79) and for promoter escape (80). We observed an accumulation of transcriptional incompetent RNA pol I devoid of RRN3 at the promoter upon IGS38 knock down, suggesting that IGS38 reduction results in an impaired recruitment of RRN3/TIF-1A and promoter escape in growing cells. The interaction of RRN3 with RNA pol I is important for its binding to the rRNA promoters and stabilises the machinery for transcription initiation (68, 73, 80, 81) but also to permit promoter escape by interacting with SL1 and DNA (79, 82). FRAP studies have shown that the RNA pol I machinery assembles as subcomplexes and RRN3 is essential in increasing the assembly efficiency and stabilising the RNA pol I at the promoter (81). Our results suggest that IGS38 favours the assembly of RRN3 competent RNA pol I and counteracts RNA pol I stalling at the 47S rRNA promoter.

The spacer-RNA and the PAPAS transcripts have both been suggested to associate with the promoter, or T0, through RNA:DNA hybrids, and subsequently recruit their interaction partners to establish a more heterochromatic chromatin state. However, the sequence of the enhancer region between the two promoters in man is different from mouse and does not harbour any strong triple helix sites (83). Recent studies in human cells have shown that R-loops resulting from transcriptional activity at the 47S rRNA promoter target silencing factors (9, 34, 35). We show that IGS38 interact with the promoter differently and we propose that IGS38 acts as a scaffold by binding directly to RRN3 and the RNA pol I transcription factor SL1. Recruitment through RNA pol I proteins is also observed in mouse ESC differentiation, in which the spacer-RNA recruits TIP-5 by binding to TTF1 (20). In human cells, cancer related ncRNAs, such as LINC01116 (TALNEC2) and SLERT, transcribed from outside the rDNA loci, are imported into the nucleolus and contribute to the hyperactive ribosomal transcription associated with malignant transformation. LINC01116 is overexpressed in a number of cancers (84) and part of the oncogenic function is performed in the nucleolus where it binds to TAF1A and TAF1D and acts as a scaffold for SL1 at the 47S rRNA promoter to enhance 47S rRNA transcription (45). We show that LINC01116 also bound directly to TAF1C, RRN3 and WSTF. It is possible that LINC01116 operates in a similar manner to IGS38 and during cancer progression it potentiates the activation of rRNA transcription by copying the function of IGS38; to stabilise the RNA machinery at the promoter to enhance transcription.

IGS38 co-localised in cross-linked chromatin with WSTF at the 47S rRNA promoter and both factors affected chromatin at the site. Knock down of IGS38 led to a less accessible promoter, encompassing the T0 site, UCE, the core element, and the TSS. The same region is also less accessible in cells in which WSTF is silenced by siRNA (31, 32). As IGS38 also binds directly to WSTF, we propose that it is involved in recruiting B-WICH to open up the promoter chromatin for transcription factors. There are differences, however, and in contrast to the IGS38 knock down, WSTF knock down does not alter the level of UBF at the promoter and the RNA pol I and SL1 are excluded (31, 32). This suggests that the IGS38 and WSTF, with their different functions co-operate to achieve a stronger UBF binding, which stabilise RNA pol I factors, such as SL1 and RRN3, at the promoter. Knock down of the IGS38 led to further chromatin alterations; a disordered CTCF boundary element with a disorganised chromatin at the spacer promoter with higher levels of H2AZ and higher levels of nucleosomes carrying histone H3K27me3 without the conventional polycomb PRC1 factor CBX2 being recruited. The histone modification H3K27me3 is also part of bivalent nucleosomes over the promoters in the poised chromatin state regulated by NuRD and CSB (21), and it is possible that IGS38 restricts the spreading of H3K27me3 and the activity of the NuRD complex at the promoters.

Our results show that the IGS38 functioned independently from the spacer-RNA and PAPAS. We only detected fragments of the spacer-RNA upstream of the TTF1 element Tsp, suggesting that the TTF1 bound to the Tsp element imposes a strong termination site in human cells. This also occurs in mouse cells, in which TTF1 bound at the Tsp forms a road block for transcription and must be removed for a read through spacer-RNA transcript to be generated and subsequently to disrupt the UBF with SL1 at the promoter (10, 13, 68). IGS38 knock down did not decrease the binding of TTF1 to the Tsp, keeping the road block. However, the binding to the T0 site was slightly reduced, but that had no effect on read through transcript. TTF1 is also involved in chromatin organisation, forming loops to activate transcription (85). Instead of altering the generation of read-through ncRNAs, IGS38 may influence the chromatin landscape and nuclear architecture by inducing RNA clouds to promote interactions (55). We did not detect any differences in PAPAS level upon IGS38 knock down and the RNAs responded differently to various stressors, suggesting that they function independently and may even operate on different rRNA gene copies. IGS38 expression was very stable and only reduced in hypotonic stressed cells, whereas PAPAS was upregulated by several stress conditions. What is regulating IGS38 is still unknown, and the response to hypertonic stress is poorly understood but the fast reaction involves a number of cellular responses to prevent cell swelling, in particular activating a number of ion channels short-term to restore normal size (86). Furthermore, IGS38 expression is stable in primary cells and cell lines, whereas PAPAS is highly expressed in differentiated cells, which have low 47S rRNA gene transcription, and it is reduced in highly proliferative cells and in malignant cells (35, 87). We postulate that IGS38 and PAPAS are functionally independent, IGS38, together with B-WICH, reinforce and maintain and active ribosomal biogenesis.

Ribosomal transcription and biogenesis are affected by many signal pathways in addition to being responsive to cellular stress. Generally, viral infections, metabolic distribution and translational stress induce nucleolar stress which activates p53 to induce apoptosis or cell cycle arrest (36, 37). In addition to p53 dependent pathways, a disrupted rRNA transcription induces an NFκB response with an accumulation of the activated transcription factors RELA and p50 in the nucleolus (37, 38). The RNA seq analysis of IGS38 knock down cells showed that the transcript was associated with an innate immune response to dsRNA; the OAS2 and CXCL11 were induced, with further ISGs, such as MDA5 from the IF1H gene, induced at low levels. OAS2, which is part of the antiviral response, is activated by dsRNA and interferon, and induces the RNAse L immune pathway to degrade RNAs and stop translation (38, 74, 75). Endogenous dsRNAs, in particular mitochondrial RNA, transposons and RNA from repetitive elements, trigger the same inflammatory response pathways as viruses (88, 89). We detected slightly higher levels of endogenous dsRNA in the cytoplasm of IGS38 knock down cells compared to control cells, but this was not sufficient to activate RNAse L. RNAse L is most potently activated by OAS3, which was not induced in IGS38 knock down cells. Overexpression of PAPAS also induces an immune response, in particular of genes in the interferon response pathway through IRF7, including OAS genes (35). Both MDA5 and IRF7 can be a response to OAS activation but also be induced independently as a response to cytoplasmic dsRNA. The coupling between rRNA transcription and an immune response may be triggered by unprocessed rRNA and dysregulated lncRNA transcription forming dsRNA from the rDNA loci. This links rRNA biogenesis to immune responses to endogenous nucleic acids and shows that dysregulation of rRNA is a potential risk to trigger sterile inflammation or autoimmunity.

**Supplementary Data are available**

## Acknowledgements - Funding

This research was funded by grants from The Swedish Cancer Society [19 0453 Pj, 22 2310 Pj]; Carl Trygger Foundation; Stockholm University to AKÖF and The Swedish Research Council [grants 2019-03853]; The Swedish Cancer Society [grant 19 0258 Pj] to NV.

We thank the Imaging Facility at Stockholm University (IFSU) for support with microscopy. Scientific illustrations were created by biorender (https://biorender.com/shortURL).

## Author Contributions

AÖF, KT, MP designed the study, AS, MP, KT, JQ, SB performed lab work. MP, KT, AÖF analysed and/or finalized the data. EE, NV performed microscopy and analysations. KT and MP performed statistical analyses. KT, AÖF, MP wrote the paper with input from AS, EE, NV.

