## Supplementary Information for "IGS38, a lncRNA from the human rDNA intergenic spacer, regulates rRNA transcription by altering rDNA chromatin organisation and activating the transcription machinery"

**Supplementary Figures and Tables – Tariq et al.**

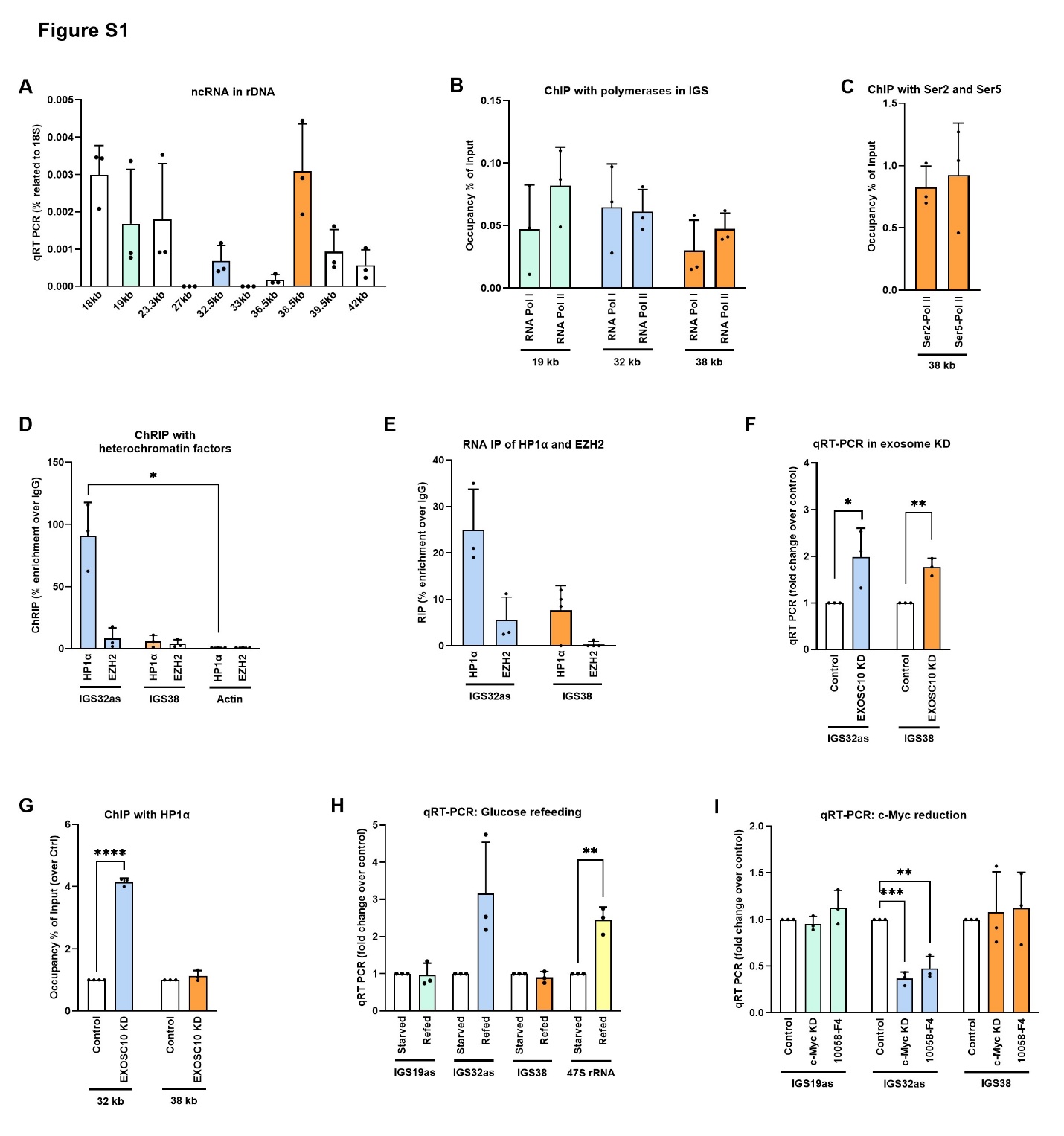

***Supplementary Figure S1*** **Noncoding RNAs from human rDNA. A**) Total RNA was extracted and region-specific primer pairs were used to amplify RNA at different position of the rDNA IGS. The position in kb from the TSS of the 47S rRNA gene is depicted below the bar. The qPCR results were normalised to 18S rRNA and are presented as % of 18S. **B**) ChIP analysis on growing HeLa cells for RNA Pol I and RNA Pol II occupancy at the positions 19 kb, 32 kb, and 38 kb from the TSS of the 47S rRNA gene. Signals are calculated as percentage of input samples. **C**) ChIP of Ser2-Pol II and Ser5-Pol II at the 38 kb position. Signals are calculated as percentage of input samples. **D**) ChRIP analysis of the association of IGS32as and IGS38 to heterochromatin using antibodies against HP1α and EZH2. RT-qPCR data is shown as percentage enrichment over IgG. **E**) RNA IP analysis to reveal direct interaction between IGS32as and IGS38 with the chromatin proteins HP1α and EZH2. RT-qPCR data is shown as percentage enrichment over IgG. **F**) IGS32as and IGS38 levels after EXOSC10 knock down. cDNA from total RNA was normalised to 18S. The results are presented as fold change over cells transfected with control siRNA **G**) ChIP-qPCR analysis of HP1α occupancy in EXOSC10 knock down cells at the positions 32 kb and the 38 kb from the TSS. Signals are calculated as percentage of input samples. **H**) qRT-PCR of ncRNA response to six-hour-glucose stimulation after starvation for 18 hours. The RNA detected is depicted under the bar. The data was normalised to 18S rRNA and is shown as fold change over control. **I**) qRT-PCR of ncRNA in response to c-MYC reduction with target siRNAs or the specific inhibitor F-10058-F4. The data was normalised to 18S rRNA and is presented as fold change over control.

All experiments present means for at least 3 biological replicates. Error bars represent standard deviation. P-values are calculated with unpaired student’s t-test: * p≤ 0.05, **p≤ 0.01,*** p≤ 0.001 and **** p≤ 0.0001.

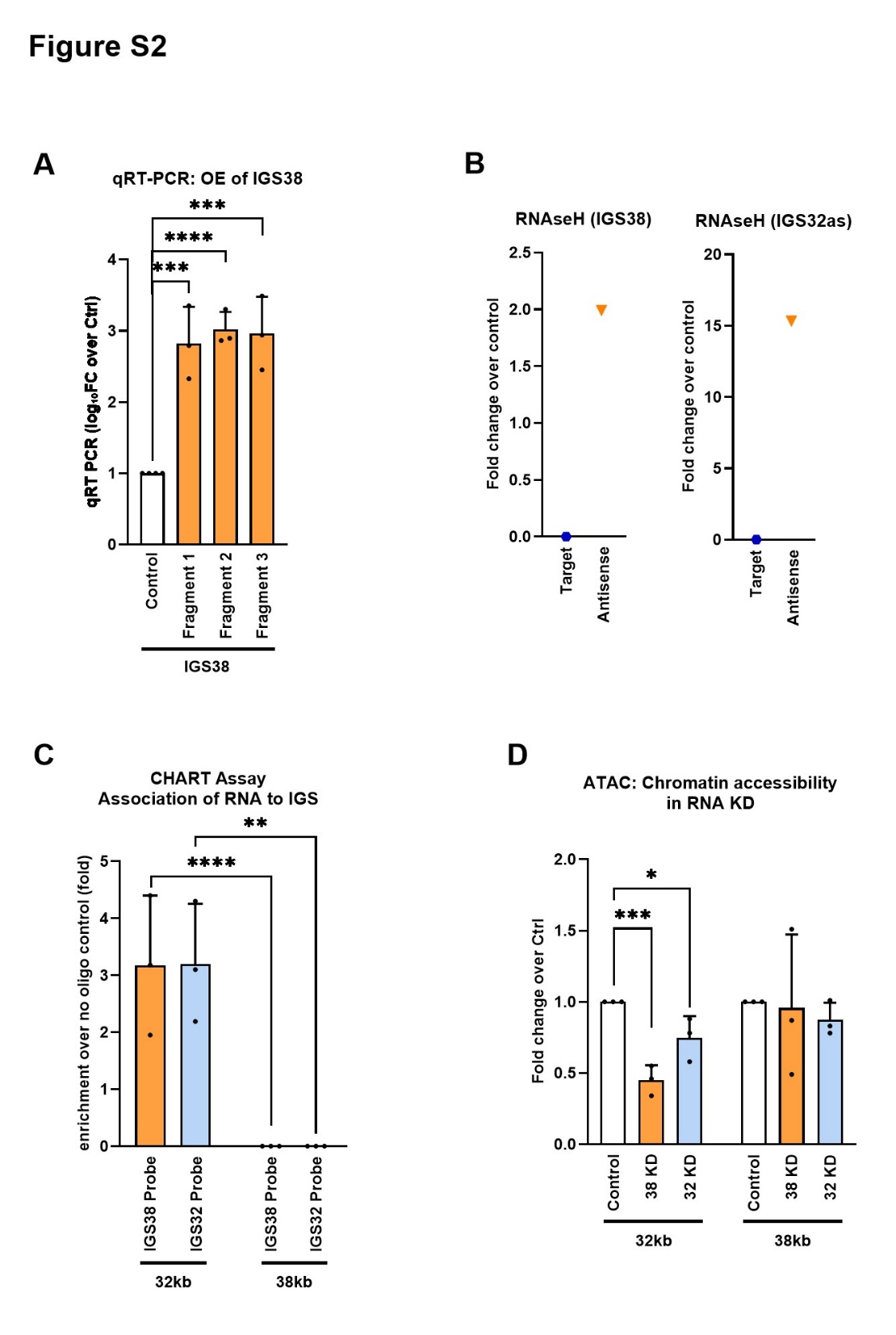

***Supplementary Figure S2*** **IGS38 interacts with the rRNA gene promoter and 32 kb region. A**) Overexpression of IGS38 from plasmids expressing three different fragments from the region¸ 38.756 bp to 39.156 bp; Fragment 1, 200 bp from position 38.759 bp to 38.957 bp; Fragment 2, 200 bp from position 38.958 kb to 39.156 kb; Fragment 3, 400 bp from 38.756 bp to 39.156 bp (Supplementary Table 1) analysed for IGS38 rRNA levels by qRT-PCR. Normalisation was done to 18S rRNA and is presented as log₁₀FC over control. **B**) RNase H treatment to identify efficient target and antisense probes. cDNA from total RNA normalised to 18S rRNA and presented as fold change over control. **C**) CHART analysis using specific probes against IGS32as and IGS38 ncRNA at the position 32 kb and 38 kb in IGS regions. Signal is presented as fold enrichment after normalising to Input signal. **D**) ATAC-qPCR analysis to assess chromatin accessibility at the 32 kb and 38 kb in the IGS regions. Signals were normalised to the signal at the 27 kb region and presented relative to control.

All experiments present means for at least 3 biological replicates. Error bars represent standard deviation. P-values are calculated with unpaired student’s t-test:* p≤ 0.05, **p≤ 0.01 and *** p≤ 0.001.

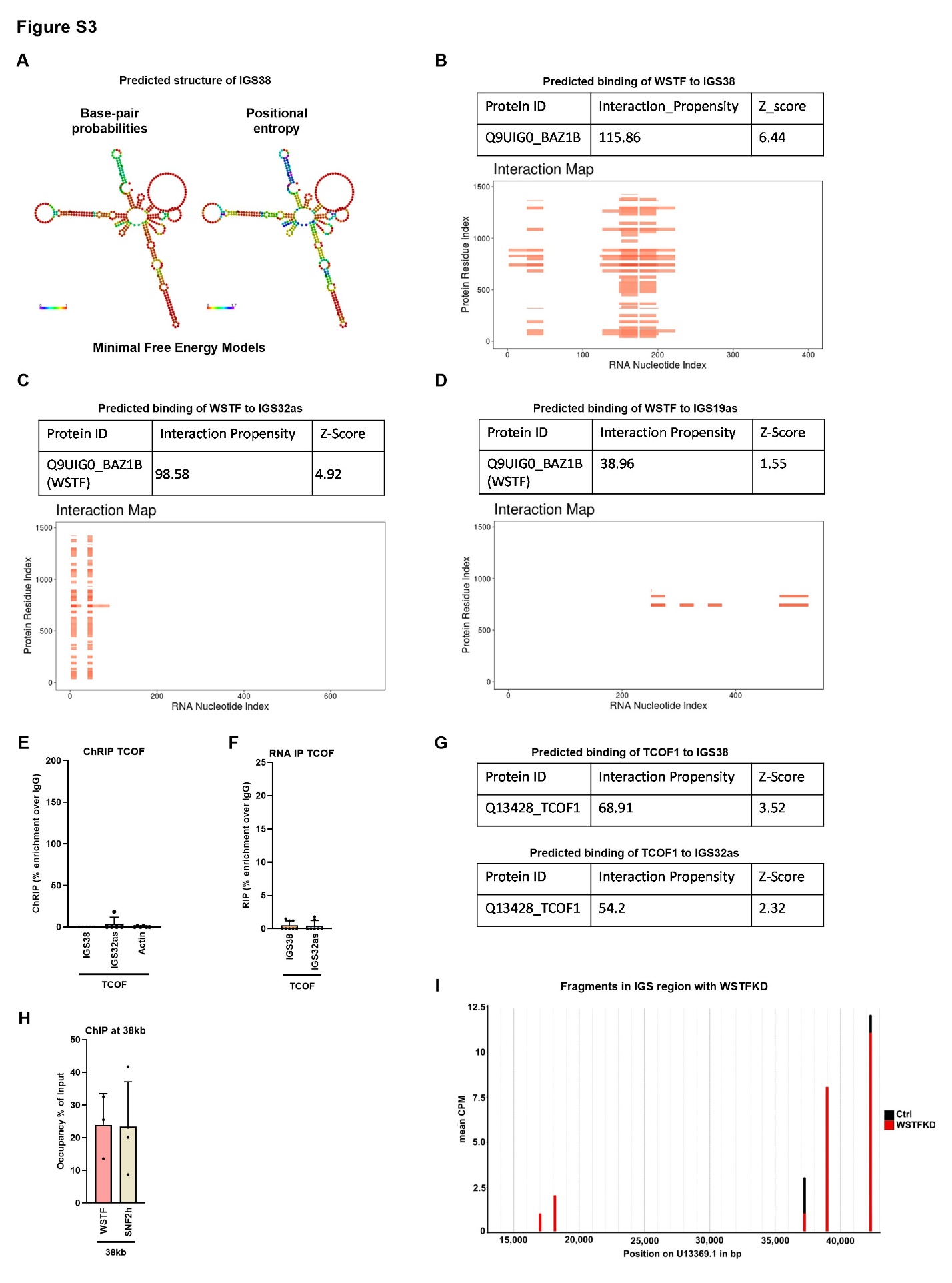

***Supplementary Figure S3*** **WSTF is predicted to bind IGS38 and both associate at the 47S rRNA promoter.** **A**) Predicted structures for IGS38 based on Maximal Free Energy with RNA Fold showing Base-pair probability in the left panel and Positional entropy in the right panel. **B**) *In-silico* catRAPID analysis of IGS38 binding to WSTF, giving the propensity score (interaction probability between one protein and one RNA) and Z-score (normalised propensity) at the top and the predicted heat map at the bottom. The varying shades of red on the heat map represent the interaction score between individual amino acid and nucleotide pair. **C**) *In-silico* catRAPID analysis of IGS32as binding to WSTF, giving the propensity score (interaction probability between one protein and one RNA) and Z-score (normalised propensity) at the top and the predicted heat map at the bottom. The varying shades of red on the heat map represent the interaction score between individual amino acid and nucleotide pair. **D**) *In-silico* catRAPID analysis of IGS19as binding to WSTF, giving the propensity score (interaction probability between one protein and one RNA) and Z-score (normalised propensity) at the top and the predicted heat map at the bottom. The varying shades of red on the heat map represent the interaction score between individual amino acid and nucleotide pair. **E**) ChRIP of the association of IGS38 and IGS32as with TCOF bound to chromatin. Actin was used as a negative control. RT-qPCR data is shown as percentage enrichment over IgG. **F**) RNA IP analysis to reveal direct interaction between IGS38 or IGS32as with TCOF. RT-qPCR data is shown as percentage enrichment over IgG. **G**) *In-silico* catRAPID analysis of IGS38 binding to TCOF (top panel) and IGS32as binding to TCOF (bottom panel), giving the propensity score (interaction probability between one protein and one RNA) and Z-score (normalised propensity). **H**) ChIP-qPCR analysis of WSTF and SNF2h occupancy at the 38 kb region. The signal is calculated as percentage of input samples. I) Fragments at different positions of the IGS, marked at the X-axis, identified in small RNA-seq in WSTF KD cells, red bars, relative to control cells, indicated in black.

All experiments present means for at least 3 biological replicates. Error bars represent standard deviation.

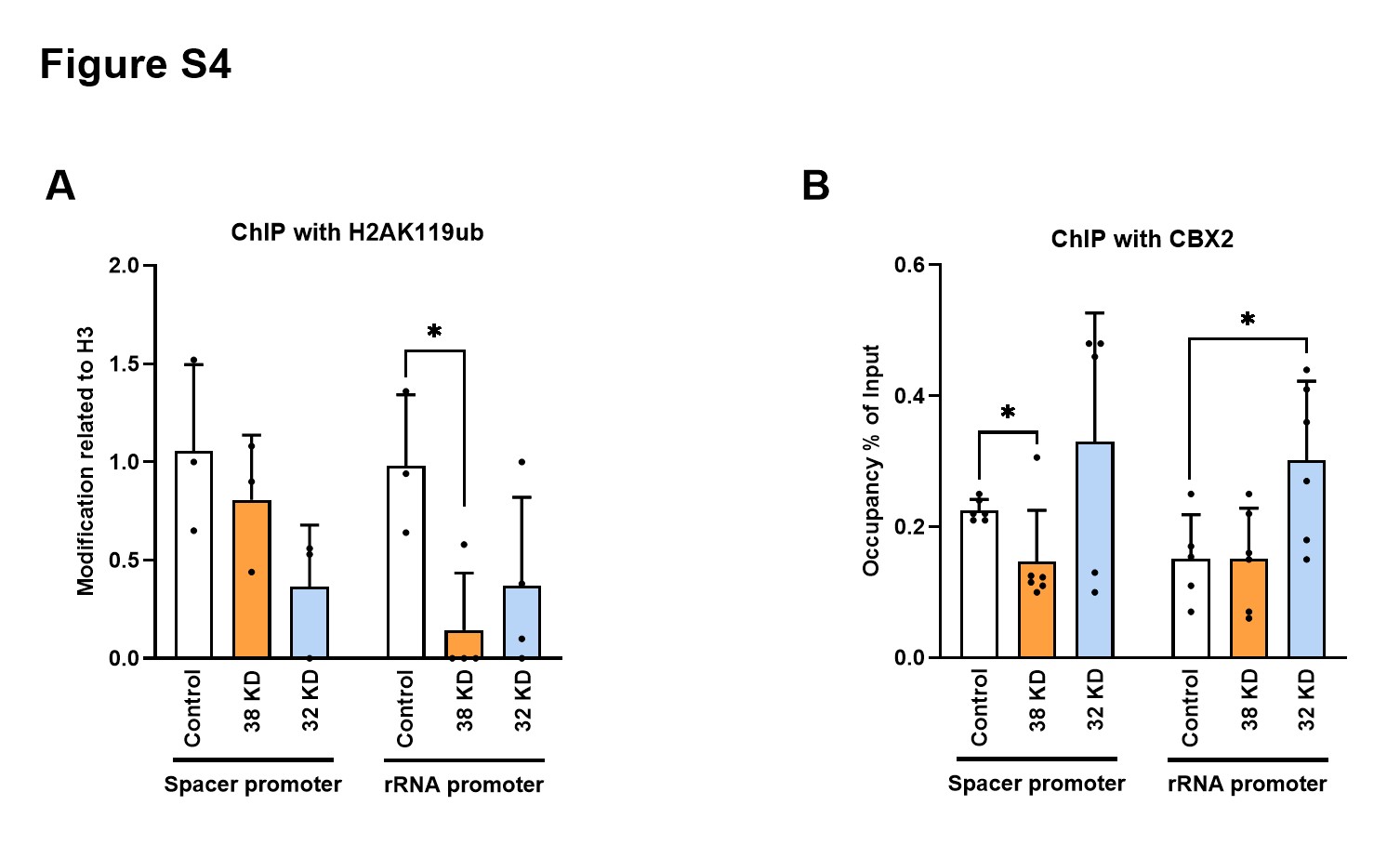

***Supplementary Figure S4*** **Chromatin landscape and transcription at the spacer promoter remain unaffected. A**) ChIP-qPCR analysis of the presence of the histone H2AK119ub at the spacer promoter and 47S rRNA promoter in control cells, IGS38 knock down cells, and IGS32as knock down cells. Signal is relative to histone H3 signal. **B**) ChIP-qPCR analysis of CBX2 occupancy at the spacer promoter and 47S rRNA promoter in control cells, IGS38 knock down cells, and IGS32as knock down cells. Signals are calculated as percentage of input samples.

All experiments present means for at least 3 biological replicates. Error bars represent standard deviation. P-values are calculated with unpaired student’s t-test:* p≤ 0.05.

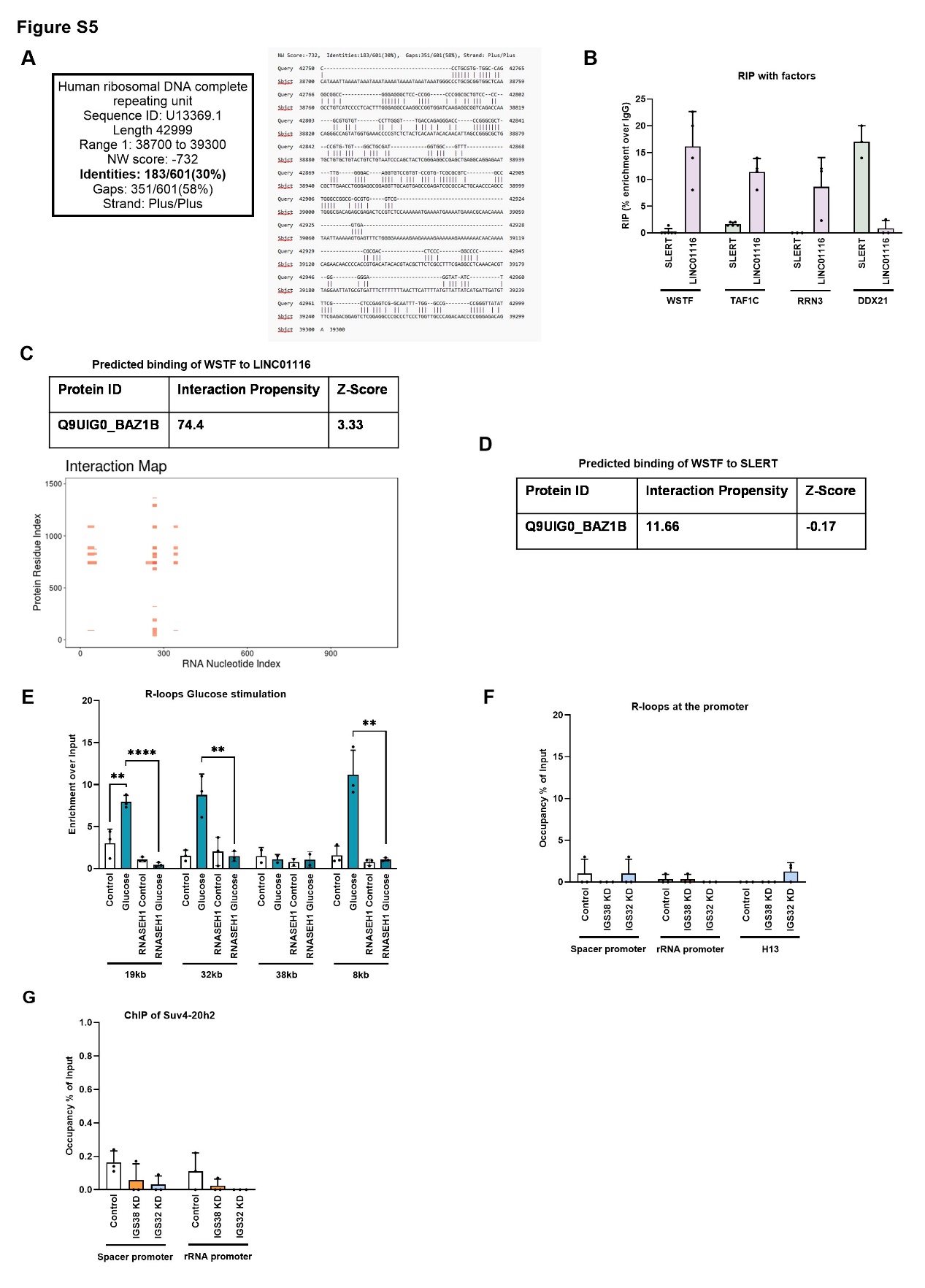

***Supplementary Figure S5****.* **No interplay exists between silencing RNA species and IGS38**.

A) Sequence homology between the 47S rRNA promoter (position 42650 to 42999 bp) and IGS38 (38700 to 39300 bp) shows only 30 % identity. **B)** RNA IP analysis to reveal interactions of the ncRNAs SLERT and LINC01116 with WSTF, TAF1C, RRN3 and DDX21. RT-qPCR data is shown as percentage enrichment over IgG. **C)** *In-silico* catRAPID analysis of LINC01116 binding to WSTF, giving the propensity score (interaction probability between one protein and one RNA) and Z-score (normalised propensity) at the top and the predicted heat map at the bottom. The varying shades of red on the heat map represent the interaction score between individual amino acid and nucleotide pair. **D)** *In-silico* catRAPID analysis of SLERT RNA binding to WSTF, giving the propensity score (interaction probability between one protein and one RNA) and Z-score (normalised propensity). **E)** DRIP-qPCR analysis of R-loop occupancy at position 19 kb, 32 kb, 38 kb and 8 kb (28S) in growing control cells, and in cells stimulated by glucose for 3 hours after 18 hours of starvation. The DRIP samples were treated with RNAse H. Signals are calculated as % of input. F) DRIP-qPCR analysis of R-loop occupancy at the spacer promoter, 47S rRNA promoter and the end of the gene at the TTS, 13 kb from the TSS, marked H13, in control cells, IGS38 knock down cells, and IGS32as knock down cells. Signals are calculated as percentage of input samples. **G**) ChIP analysis of Suv4-20h2 occupancy at the spacer promoter and 47S rRNA promoter in control cells, IGS38 knock down cells, and IGS32as knock down cells. Signals are calculated as percentage of input samples.

All experiments present means for at least three biological replicates (except for 38kb in E, which is based on two replicates). Error bars represent standard deviation. P-values are calculated with unpaired student’s t-test: * p≤ 0.05 and **p≤ 0.01.

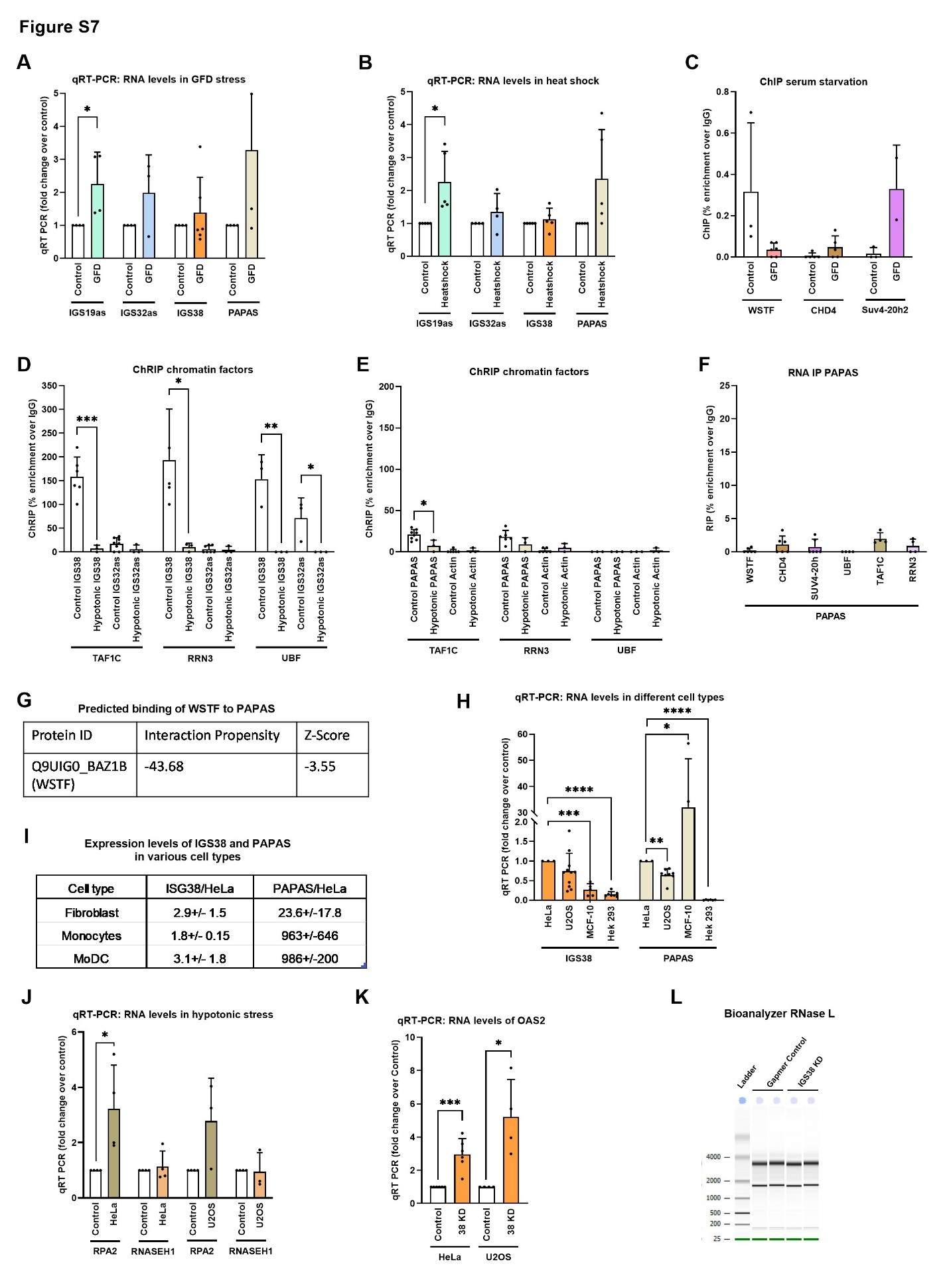

***Supplementary Figure S6.* ncRNAs in response to different kind of stress and IGS38 expression in primary cells and cancer cell lines. A**) qRT-PCR of the RNA levels of IGS19as, IGS32as, IGS38, and PAPAS in cells exposed to growth factor depletion (GFD) for 48 hours and control cells. The data was normalised to 18S rRNA and shown as fold change over control. **B**) qRT-PCR of the RNA levels of IGS19as, IGS32as, IGS38, and PAPAS in cells exposed to heat shock stressed and control cells. The data was normalised to 18S rRNA and shown as fold change over control. **C**) ChIP-qPCR analysis of WSTF, CHD4 and Suv4-20h2 occupancies in control and GFD stressed cells. The data is shown as percentage enrichment over IgG. **D**) ChRIP of the association of IGS38 and IGS32as with TAF1C, RRN3 and UBF bound chromatin in cells exposed to hypotonic stress. qRT-PCR data is shown as percentage enrichment over IgG. **E**) ChRIP of association of PAPAS with TAF1C, RRN3 and UBF bound chromatin. Associated actin was used as a control. qRT-PCR data is shown as percentage enrichment over IgG. **F**) RNA IP to analyse direct interactions between PAPAS and the factors WSTF, CHD4, Suv-4h2, UBF, TAF1C, and RRN3 as marked under the bars. qRT-PCR data is shown as percentage enrichment over IgG. **G**) *In-silico* catRAPID analysis of WSTF binding to PAPAS, giving the propensity score (interaction probability between one protein and one RNA) and Z-score (normalised propensity). **H**) qRT-PCR to analyse RNA levels of IGS38 and PAPAS in different cell lines; HeLA cells, U2OS cells, MCF10 cells, and HEK 293 cells. The data is normalised to 18S rRNA and presented as fold related to the level in HeLa cells. **I**) Table of expression levels of IGS38 and PAPAS related to the level in HeLa cells in human fibroblasts, human monocytes and monocyte derived dendritic cells (moDCs). **J**) qRT-PCR to analyse RNA levels of RPA2 and RNASEH1 in HeLa and U2OS cells in cells exposed to hypotonic stress and control cells. The RNA is normalised to 18S rRNA and shown as fold change over control. **K**) qRT-PCR analysis of OAS2 in IGS38 knock down HeLa cells and U2OS cells related to control cells transfected with LNA-DNA Gapmer. The data is normalised to 18S rRNA and presented as fold change over control. **L**) Bioanalyzer analysis to assess RNaseL activity in IGS38 knock down cells and Gapmer control cells. The bp marker is in the lane marked Ladder, and the sizes are indicated to the left.

All experiments present means for at least three biological replicates. Error bars represent standard deviation. P-values are calculated with unpaired student’s t-test: * p≤ 0.05, **p≤ 0.01, *** p≤ 0.001 and **** p≤ 0.0001.

**Supplementary Table 1**

| **RNA expression primers (initial screening) F R** | | |
| --- | --- | --- |
| 18kb | 5’-GTTGACGTACAGGGTGGACTG-3’ | 5’-GGAAGTTGTCTTCACGCCTGA-3’ |
| 19kb | 5’-CGTCGTAACTCACTCCTGC-3’ | 5’-GCAGAGGTTGCGGTGAGCCA-3’ |
| 23.3kb | 5’-TTTAAAAGCGCGCGGCCC-3’ | 5’-CCTATGGCAGAGAGGACACG-3’ |
| 27kb | 5’-GACATTTGCAGGCAGGCATC-3’ | 5’-GGCCTCCAGAAGGGAGAGA-3’ |
| 32.5kb | 5’-GGAGTGCGATGGTGTGATCT-3’ | 5’-TAAAGATTAGCTGGGCGTGG-3’ |
| 36.5kb | 5’-GGGATTACAGGGAGGAGC-3’ | 5’-GTAGCCGGCCAGTCACG-3’ |
| 38.5kb | 5’-CTTTGGGAGGCCAAGGCC-3’ | 5’-GATCTCGGCTCACTGCAAC-3’ |
| 39.5kb | 5’-CTGATTCTCCCCAGATGTAG-3’ | 5’-CAAGAGTGAAACTCTGTCTC-3’ |
| 42kb | 5’-AGAGGGGCTGCGTTTTCGGCC-3’ | 5’-CGAGACAGATCCGGCTGGCAG-3’ |
| **RNA expression primers F R** | | |
| IGS19as | 5’-CGTCGTAACTCACTCCTGC-3’ | 5’-GCAGAGGTTGCGGTGAGCCA-3’ |
| IGS32as | 5’-CCTAGTAAAAGCTGGCCGATC-3’ | 5’-CCCAAGTCTGGTTGATCCTTG-3’ |
| IGS38-1 | 5’-GAGTTTCTGGGGAAAAAGAAG-3’ | 5’-GAGAAGCGTACGTGTATGTCAC-3’ |
| IGS38-2 | 5’-GTGACATACACGTACGCTTCTC-3’ | 5’-GCCTCCGAGACTCCGTC-3’ |
| 47S rRNA | 5’-CTCCGTTATGGTAGCGCTGC-3’ | 5’-GCGGAACCCTCGCTTCTC-3’ |
| 18S rRNA | 5’-CGACGACCCATTCGAACGTCT-3’ | 5’-CTCTCCGGAATCGAACCCTGA-3’ |
| Actin mRNA | 5’-GGACTTCGAGCAAGAGATGG-3’ | 5’-AGCACTGTGTTGGCGTAC-3’ |
| SLERT | 5’-TTAGTCAGCTCAGGCCCAGT-3’ | 5’-AAGTGCTCCACCAACTCCAG-3’ |
| LNC01116 | 5’-GTTCAAGTGCGTCCGGGTTT-3’ | 5’-CGGACTTCTTTTCCAGGCGG-3’ |
| 42235 | 5’-GGTGGCGCCAGAGCTGTG-3’ | 5’-GAAGCGCAGCGACAGCCTC-3’ |
| 42370 | 5’-GTAGCTCCCGAGGCCCG-3’ | 5’-CTCGGAAACGAATTCGGCC-3’ |
| 42807 | 5’-GTGTCCTGGGGTTGACCAG-3’ | 5’-CGACTCGGAGCGAAAGATA-3’ |
| PAPAS (-49) | 5’-GGTATATCTTTCGCTCCGAG-3’ | 5’-GACGACAGGTCGCCAGAGGA-3’ |
| PAPAS (-145) | 5’-GCGATGGTGGCGTTTTTGG-3’ | 5’-GACGACAGGTCGCCAGAGGA-3’ |
| **DNA primers for ChIP/ATAC/CHART F R** | | |
| Spacer region 42235 (CHIP) | Same as spacer region 42235 above | |
| Spacer region 42281 | 5’-CGTGCAGGTTTATGTGGG-3’ | 5’-CGGGCCTCGGGAGCTACG-3’ |
| Spacer region 42370 | Same as spacer region 42370 above | |
| 42453 | 5’-CGCTCATCCTGGCCGTC-3’ | 5’-GAGACGGCGCTAGGAAAGAC-3’ |
| 42589 | 5’-GATCCTTTCTGGCGAGTCC-3’ | 5’-GGCTTTTACGAAGGCCGAG-3’ |
| 42760 | 5’-CGTGGATTCCGGAAGAGCC-3’ | 5’-GGAGGGACGAAGGCTCTC-3’ |
| 42808 | 5’-GTCCTTGGGTTGACCAGAG-3’ | 5’-GTCCACAGGCACAGGCACAG-3’ |
| 2c | 5’-CTGACACGCTGTAATCTGG-3’ | 5’-GGCGCGAGGACGGAACTC-3’ |
| 85c | 5’-CTAGCCGGCCGCGCTCC-3’ | 5’-CCTTCCGCCGCTCCCGG-3’ |
| ETS | 5’-GGCGGTTTGAGTGAGACGAGA-3’ | 5’-ACGTGCGCTCACCGAGAGCAG-3’ |
| 4kb | Same as 18S rRNA | |
| 8kb | 5’-AGTCGGGTTGCTTGGGAATGC-3’ | 5’-CCCTTACGGTACTTGTTGACT-3’ |
| 47SP | 5’-GCGATGGTGGCGTTTTTGG-3’ | 5’-AGCGACAGGTCGCCAGAGGA-3’ |
| IGS32kb | Same as expression primer IGS32as | |
| IGS38kb | Same as expression primer IGS38-2 | |
| IGS27kb | 5’-CCTTCCACGAGAGTGAGAAGCG-3’ | 5’-CTCGACCTCCCGAAATCGTAC-3’ |
| ActinP | 5’-5′-CAGAAGGATTCCTATGTGGG, | 5’- TGGATAGCAACGTACATGGC |
| **LNA-DNA Gapmers for knockdown (Qiagen)** | | |
| Control | 5’-AACACGTCTATACGC-3’ |  |
| IGS32as 1-2 | 5’-TAACAACACAAGGATC-3’ | 5’-ATCGGAAGAGAAGGCA-3’ |
| IGS38 1-2 | 5’-CGCATAATTCCTAACG-3’ | 5’-TGTGAGTAGAGACGG-3’ |
| **siRNA** | | |
| Control | 5’-GUGCGAGGGGGUUGUAAUCTT-3’ | |
| WSTF | 5’-GGAAGGAGAGAGAGUAUUA-3’ | 5’-CUAAGGCACUUCACUUAGA-3’ |
| EXOSC10 | 5’-GCUGCAGCAGAACAGGCCAUU-3’ | |
| **Probes (Northern Blot)** | | |
| IGS19 RNA | 5’-GACTCGAGCGATCCTTCCACCTCAGCCTCCAGAGTACAGAGCCTGGGACC-3’ | |
| IGS32 RNA | 5’-TCTTGAGACGCGTGTCGCTCTGTCGCCCAGGCTGGAGTGCGATGGTGTGA-3’ | |
| IGS38 RNA | 5’-GCTAATGTTGTGTATTGTGAGTAGAGACGGGGTTTCACCATACTGGCCCT-3’ | |
| **Probes (CHART) Target Probe Antisense** | | |
| 32P labelled | 5’-TCTTGAGACGCGTGTCGCTCTGTCGCCCAGGCTGGAGTGCGATGGTGTGA**[SpC18][BIOTEG]**-3’ | 5’-TCACACCATCGCACTCCAGCCTGGGCGACAGAGCGACACGCGTCTCAAGA-**[SpC18][BIOTEG]**-3’ |
| 38P labelled | 5’-TGTATTGTGAGTAGAGACGG**[SpC18][BIOTEG]**-3’ | 5’-CCGTCTCTACTCACAATACA**[SpC18][BIOTEG]**-3’ |
| **Probes (FISH)** | | |
| 38-1 | 5’-**[DIG]**CGAGAAGCGTACGTGTATGT**[DIG]**-3’ | |
| 38-2 | 5’-**[DIG]**AGGCCTCGAAAGGCGAGAAG**[DIG]**-3’ | |
| 47S | 5’-**[DIG]**CCGTCGCGGCTCGGACCCGGCCCGGGAGAGCACGACGTCACCACATCGAT[**DIG]**-3’ | |
| Bacterial | 5’-**[DIG]**CCAGGGTCAACAACGCGACGGTAACCGTCA**[DIG]**-3’ | |
| **Sequences for Overexpression of IGS38 with snoRNA boxes** Box C **and** Box D | | |
| Fragment 1 | 5’-TGCAATGATGTCGTAATTTGCGTCAGCCTGTCATCCCCTCACTTTGGGAGGCCAAGGCCGGTGGATCAAGAGGCGGTCAGACCAACAGGGCCAGTATGGTGAAACCCCGTCTCTACTCACAATACACAACATTAGCCGGGCGCTGTGCTGTGCTGTACTGTCTGTAATCCCAGCTACTCGGGAGGCCGAGCTGAGGCAGGAGAATCGCTTGAACCTGGGAGGCGGAGGTTGCAGTGAGCCGAGATCGCGCCACTGCAACCCAGCCTGGGCGACAGAGCGAGACTCCGTCTCCAAAAAATGAAAATGAAAATGAAACGCAACAAAATAATTAAAAAGTGAGTTTCTGGGGAAAAAGAAGAAAAGAAAAAAGAAAAAAACAACAAAACAGAACAACCCCACCGTGACATACACGTACGCTTCTCGACGAAAATTCTTACTGAGCAA-3’ | |
| Fragment 2 | 5’-TGCAATGATGTCGTAATTTGCGTCAGCCTGTCATCCCCTCACTTTGGGAGGCCAAGGCCGGTGGATCAAGAGGCGGTCAGACCAACAGGGCCAGTATGGTGAAACCCCGTCTCTACTCACAATACACAACATTAGCCGGGCGCTGTGCTGTGCTGTACTGTCTGTAATCCCAGCTACTCGGGAGGCCGAGCTGAGGCAGGAGAATCGCTTGAACCTGGGAGGCGACGAAAATTCTTACTGAGCAA-3’ | |
| Fragment 3 | 5’-TGCAATGATGTCGTAATTTGCGTCGGAGGTTGCAGTGAGCCGAGATCGCGCCACTGCAACCCAGCCTGGGCGACAGAGCGAGACTCCGTCTCCAAAAAATGAAAATGAAAATGAAACGCAACAAAATAATTAAAAAGTGAGTTTCTGGGGAAAAAGAAGAAAAGAAAAAAGAAAAAAACAACAAAACAGAACAACCCCACCGTGACATACACGTACGCTTCTCGACGAAAATTCTTACTGAGCAA-3’ | |

**Supplementary Table 2**

| **Antibody** | **Company/Cat. Number** |
| --- | --- |
| CBX2 | Abcam, ab80044 |
| CHD4 | Abcam, ab72418 |
| c-MYC | Abcam, ab32072 |
| CSB | Abcam, ab66598 |
| CTCF | Abcam, ab70303 |
| DDX21 | Abcam, ab182156 |
| EZH2 | Abcam, ab3748 |
| H2Aub119 | Abcam, ab193203 |
| H2Az | Abcam, ab188314 |
| H3 | Abcam, ab1791 |
| H3K27Ac | Abcam, ab4729 |
| H3K27me3 | Abcam, ab6002 |
| H3K9me3 | Abcam, ab8898 |
| HP1α | Abcam, ab109028 |
| IgG Control | Abcam, ab18443 |
| J2 | Cell Signaling, 76651 |
| RNA Pol I | Aviva Systems Biology, OASG06470 |
| RNA Pol II | Abcam, ab26721 |
| RRN3 | Abcam, ab112052 |
| Ser2 | Abcam, ab5095 |
| Ser5 | Abcam, ab5131 |
| Suv4-20h2 | Abcam, ab104875 |
| S9.6 | Kerafast, ENH001 |
| TAF1C | Abcam, ab234957 & ab178690 |
| WSTF | Abcam, ab70263 & Cell Signaling 2152 |
| TBP | Abcam, ab51841 |
| TCOF | Abcam, ab65212 |
| TTF1 | GeneTex, GTX129804 |
| UBF | Abcam, ab75781 |
